# Optimized pipeline for generating highly potent neutralizing Affitins: application to SARS-CoV-2 spike protein

**DOI:** 10.1101/2025.04.03.646589

**Authors:** Thomas Maurice, Mélanie Laporte, Nicolas Barquilla, Agnès Fortun, Klara Echasserieau, Florence Guivel-Benhassine, Arménio Barbosa, Pauline Cossard, Cyril Planchais, François Anna, Pierre Charneau, Hugo Mouquet, Cécilia Roque, Olivier Schwartz, Barbara Mouratou, François Davodeau, Karine Bernardeau, Blandine Monel, Frédéric Pecorari

## Abstract

Viral infections represent a global health threat, causing three million deaths annually. Antiviral strategies primarily target viral genome replication or prevent virus entry into host cells. While most FDA-approved antiviral drugs are small chemical molecules, protein-based therapies like monoclonal antibodies (mAbs) offer advantages such as broad and high neutralizing activities, and ability to recruit immune responses to enhance viral clearance. Additionally, alternative protein scaffolds with favorable properties have been developed to substitute or complement mAbs. We have developed Affitins, a novel class of small artificial affinity proteins (7 kDa) derived from hyperthermophilic archaea. Affitins, selected from high-diversity libraries (∼10^12^ variants) against any target protein, are highly stable, easy to engineer, and cost-effective to produce.

Here, we developed a pipeline to generate Affitins in different multimerization formats targeting the receptor-binding domain (RBD) of the SARS-CoV-2 spike protein through ribosome display selection and assessed their efficacy. The most potent candidates, dimerized or trimerized into 17–27 kDa proteins, displayed high thermal stability, strong binding affinities, and potent neutralization against various SARS-CoV-2 variants, with IC_50_ values as low as 32 pM. We also report the first hexameric Affitins, formed by dimerizing trimers through Fc fragment fusion (∼106 kDa), achieving potent neutralization with an IC_50_ of 0.8 pM, ranking them among the most potent B1.351 SARS-Cov-2 neutralizing proteins. Our findings highlight Affitins as promising antiviral agents, demonstrating their versatility for engineering affinity proteins with higher valencies than that of antibodies while retaining a lower molecular weight, thereby expanding the toolkit for combating viral threats.

## 1. Introduction

Most FDA-approved antiviral drugs are small chemical molecules [1]. However, this class of molecules can exhibit off-target toxicity, leading to undesirable side effects. Additionally, they have a short half-life, necessitating frequent dosing, and possess a limited capacity to engage the host immune system. Although, protein-based therapies, including monoclonal antibodies (mAbs), are less common, they have become an attractive antiviral option due to their high specificity, reducing the likelihood of adverse side effects. They also generally exhibit longer half-lives than small chemical molecules. Monoclonal antibodies have been approved by the FDA for various viral infections. For example, Palivizumab is used to prevent Respiratory Syncytial Virus (RSV) infections in high-risk infants by targeting the F protein, blocking viral entry [2]. Ibalizumab is approved for treating multi-drug-resistant HIV-1, preventing viral entry by binding to the CD4 receptor on T-cells [3]. When used in combination, mAb “cocktails” (ie with different binding specificities) can reduce or delay the risk of viral escape, as they require the virus to mutate at multiple binding sites simultaneously [4]. For instance, REGN-EB3, a cocktail of three mAbs, treats Ebola by binding different sites on the viral glycoprotein, providing robust neutralization [5]. Beyond neutralization, mAbs can recruit other immune components to help clear infections, utilizing mechanisms such as antibody-dependent cellular cytotoxicity (ADCC) or complement activation [6–8].

However, the large size of 150 kDa of mAbs can hinder access to deep grooves on viral surfaces that are often less prone to escape mutations making them good targets [9,10]. Additionally, repeated epitopes are often displayed on the surface of viruses. While mAbs are naturally bivalent, they are not well-suited for designing multimeric proteins with enhanced avidity to fully exploit these repeated antigenic structures. To address these drawbacks, non-immunoglobulin protein scaffolds have been developed during the last three decades (see [11] for a review). Derived from proteins with diverse biological functions, these affinity scaffolds are evolved *in vitro*, by coupling combinatorial mutagenesis and selections, to recognize desired targets. They offer several advantages, including a low molecular weight (<15 kDa), ease of engineering into multivalent formats with increase in vivo half-life, optional capability to engage the host immune system, high thermal and structural stability, and low production costs.

In this work we explored the use of Affitins as a scaffold protein to neutralize severe acute respiratory syndrome coronavirus 2 (SARS-CoV-2). Affitins are small affinity proteins (7 kDa) originating from the Suld7d protein family, found in hyperthermophilic archaea *Sulfolobus* [12]. Through the randomization of 10 to 14 residues within the DNA-binding site of Sac7d, Affitins in their monomeric form have been characterized to specifically bind a wide range of targets, such as human, animal and bacterial proteins, and living bacteria [13–16]. We further demonstrated that multimerizing Affitins via dendrimers is an effective strategy to enhance avidity-driven binding to bacteria [17]. Affitins composed of a unique polypeptide chain, lack cysteine, and exhibit high stability under diverse conditions [13,15,16,18], positioning them as a viable substitute or complement to antibodies.

SARS-CoV-2, the virus responsible for COVID-19, initiates infection by binding to the host receptor, angiotensin-converting enzyme 2 (ACE2), via the receptor-binding domain (RBD) of its spike (S) glycoprotein, anchoring viral particles to the cell surface. In response, several monoclonal antibodies (mAbs) targeting the RBD have been developed, with some approved for clinical use, to effectively neutralize the virus [19]. As the virus continues to evolve with new variants, often carrying mutations in the RBD, efficacy of existing mAbs, including those approved by FDA, is reducing with some exceptions [20,21]. Other emerging viruses pose significant risks to global health because of their potential for rapid transmission, high mortality rates, and capacity to evade existing medical countermeasures including small chemical molecules [22]. Expanding our arsenal of antiviral agents with diverse mechanisms can enhance our ability to combat both current and future viral threats. Consequently, it is crucial to validate novel tools with distinct properties from traditional small chemical molecules and mAbs that can effectively neutralize viruses.

We identified novel Affitins that bind to the SARS-CoV-2 RBD using ribosome display selections and evaluated different formats and levels of Affitin multimerization. The most effective candidates, dimerized or trimerized into proteins of 17 to 27 kDa, demonstrated high thermal stability, strong binding affinities, and neutralization capability against multiple SARS-CoV-2 variants, with IC_50_ values as low as 32 pM. Additionally, this work describes the first hexameric Affitins, obtained by dimerizing trimers with a Fc fragment fusion, demonstrating potent neutralization with an IC_50_ of 0.8 pM. These findings highlight the potential of Affitins as versatile antiviral agents and enabled us to establish a pipeline for their generation.

## 2. Materials and methods

### 2.1. Protein production

The protein sequences produced for this study are shown in the Table S1. After purification, the homogeneity and size of proteins were evaluated by sodium dodecyl sulfate-polyacrylamide gel electrophoresis (SDS-PAGE), by exclusion chromatography on Superdex 75 or 200 10/300 column (Cytiva) or mass photometry (Two MP, Refeyn). The concentrations of proteins were determined by measurement of the optical density at 280 nm (molar extinction coefficients of proteins are indicated in Table S1). All protein solutions were filtered through a 0.22 µm membrane, aliquoted, and stored in PBS (pH 7.4) at -80°C.

#### 2.1.1. Production and purification of proteins in mammalian cells

Mammalian expression vectors pCAGGS-spike and pCAGGS-RBD including an hexahistidine tag in C-terminal were kindly supplied by Florian Krammer [23] and Sebastiano Pasqualato. The encoding sequences of both vectors were modified by introduction of mutations K417N, E484K and N501Y and an Avitag was added in C-terminal (Table S1). Codon optimization for human expression, gene synthesis and cloning were performed by Genecust. Mammalian expression vector pTwist-ACE2-Fc mouse IgG2a including a histidine tag in C-terminal were kindly supplied by Kelvin Lau.

These three proteins were produced in Expi293F cells (ThermoFisher Scientific) by transient transfection using Expifectamine 293 transfection kit (Thermo Fisher Scientific, A14635). Supernatant from transfected cells were harvested on day 4 post-transfection by centrifugation for RBD and spike, on day 5 post-transfection for ACE2. Each supernatant was dialyzed in equilibration buffer for further purification.

Chromatography was made on NGC medium-pressure systems from Bio-Rad or an FPLC system from Cytiva. spike protein was purified on Histrap column (Cytiva) at 4°C in 50 mM phosphate pH 8.0 buffer containing 0.5 M of NaCl and 5 mM of imidazole for column equilibration and washing. Elution was performed with 50 mM phosphate pH 8.0 buffer supplemented with 0.5 M of NaCl and 0.5 M of imidazole. Purified protein was then dialyzed in 50 mM phosphate pH8 buffer with 150 mM of NaCl.

RBD protein was purified on Histrap column (Cytiva) in PBS pH 7.4 buffer containing 0.5 M of NaCl and 25 mM of imidazole for column equilibration and washing. Elution was performed with PBS pH 7.4 buffer supplemented with 0.5 M of NaCl and 0.5 M of imidazole. RBD eluted proteins were dialyzed against PBS pH 7.4. A part of RBD protein was biotinylated with the BirA enzyme (Avidity, Denver, CO) for 5 h at 30°C, desalted on Hiprep 26/10 Desalting column (GE, Healthcare) and dialyzed overnight à 4°C in PBS.

ACE2 supernatant from transfected Expi293 F cells was dialyzed against 20 mM phosphate pH 7 buffer. Purification by Protein G column (Histrap, Cytiva) was carried-out at 4°C and 0.1 M citrate pH 3.0 buffer was used for elution step neutralized with 1 M Tris-HCl buffer pH 9 in elution fractions. Purified ACE2 protein was then dialyzed in PBS buffer.

The Affitins-Fc fusions were produced from Expi293 F cells, and were firstly purified using a Protein A column (Cytiva) using PBS pH 7.4 as running buffer and 0.1 M citrate buffer pH 3.3 for elution. After neutralization of the collected fractions by 1 M Tris-HCl buffer pH 9.0, these fusions were further purified by size-exclusion chromatography on a Superdex 200 column (Cytiva).

Finally, proteins were snap frozen in liquid nitrogen and stored at -80°C. Proteins were analyzed by SDS-Page and by size exclusion chromatography on Superdex 75 or 200 for RBD and ACE2 proteins respectively. The interaction between RBD or spike protein with the receptor ACE2 was validated by ELISA, NanoDSF (data not shown) and SPR (Figure S1).

#### 2.1.2. Production of proteins in bacteria

All Affitins were expressed in the cytoplasm of *Escherichia coli* (*E. coli)*. All synthetic DNA sequences coding for monomeric Affitins were supplied by GeneCust. These synthetic sequences were inserted via BamHI and HindIII restriction sites in the pFP1001 plasmid [24], a derivate of the pQe30 vector (Qiagen) encoding proteins fused to a RGS-hexa histidine tag at their N-terminal [24]. These plasmids were used to transform the *E. coli* DH5α I^q^ strain (Thermo Fisher Scientific) and the Affitins were expressed and purified as described previously [25] using a Ni-NTA resin (Cytiva) and size-exclusion chromatography on a Superdex 75 gel filtration column (Cytiva) equilibrated with PBS pH 7.4.

The multimeric Affitins were built by connecting Affitins with a linker encoding a 20-amino-acid segment of human muscle aldolase (HMA) [26]. The corresponding DNA sequences were generated using the GoldenGate approach [27]. The destination vector pFP1001-GG was created from pFP1001 in which the BsaI restriction site was silenced by PCR using primers GG-BsaI-del-F and GG-BsaI-del-R (see Table S2 for primer sequences). This vector was amplified and linearized by PCR using the primer pairs GG-1001-2-F/GG-1001-R and GG-1001-3-F/GG-1001-R. The resulting PCR products were then utilized, after digestion for 2h at 37°C with DpnI (5U) and ExoI (10U) enzymes (Thermo Fisher Scientific), for subsequent GoldenGate assemblies to construct dimeric and trimeric Affitins, respectively. In order to address Affitin and HMA linker fragment to a given position in the multimer, each corresponding Affitin or HMA DNA sequence subcloned in pFP1001 was submitted to a PCR using oligonucleotides GG-AFN.1-F/GG-AFN.1-R, GG-AFN.2-F/GG-AFN.2-R, GG-AFN.3-F/GG-AFN.3-R, GG-HMA.1-F/GG-HMA.2, GG-HMA.1-F/GG-HMA.1-R to create specific cohesive 5’ and 3’ends. Approximately 9 ng of each DNA encoding Affitins, 4 ng of each DNA HMA and 75 ng of linearized pFP1001-GG were mixed with 15 U of BsaI-HFv2 (New England BioLabs) and 2.5 U of T4 DNA ligase (New England BioLabs) under T4 DNA ligase buffer conditions followed by the addition of dH_2_O to a volume of 20 μL. The mixture was incubated in a thermocycler for 60 cycles of 37°C for 1 min and 16°C for 1 min, and then incubated at 60°C for 5 min for deactivation. Ten microliters of this mixture were used for transformation of *E. coli* DH5α I^q^ competent cells (Thermo Fisher Scientific). After 1 h incubation at 37°C, the cells were plated on LB agar containing Amp and incubated overnight at 37°C. Multimeric assemblies were controlled by picking isolated colonies, purification of plasmidic DNA using Wizard Plus SV Minipreps DNA Purification kit (Promega) and Sanger sequencing (Eurofins). The multimeric Affitins were then produced as described for monomeric Affitins.

### 2.2. Selection of RBD-specific Affitins by ribosome display

Three rounds of selection were carried-out using the Affitin libraries L5 and L6 [28]. For the rounds of selection 1 and 2, the spike protein (10 µg/mL in PBS) was immobilized on a Maxisorp plate (Thermo Fisher Scientific) overnight at 4°C. The selection wells were then blocked for 1h at RT with PBS containing 0.5 % bovine serum albumin (BSA). The wells were washed three times with PBS, and once with washing buffer (WBT, 50 mM Tris-acetate pH 7.4, 150 mM NaCl, 50 mM Magnesium acetate, 0.1% (v/v) Tween-20). Ribosome display selection with the translation mix containing mRNA-ribosome-Affitin complexes was performed at 4 °C as previously described [28]. Briefly, the translation mix was added to the wells with the immobilized target and incubated for 1h with gentle shaking. The unbound complexes were removed by washing with WBT containing 0.5% BSA and the mRNA were collected with elution buffer (50 Mm Tris-acetate pH 7.4, 150 mM NaCl, 20 mM ethylenediaminetetraacetic acid (EDTA)). The round 3 was performed in solution using 5 nM of biotinylated RBD. The translation mix was initially pre-incubated with 25 µL of streptavidin-coated magnetic beads (M270, Thermo Fisher Scientific) for 15 minutes at 4 °C to eliminate potential bead binders. The mixture was then transferred to a new tube and incubated with biotinylated RBD at 5 nM for 1h at 4°C. The complexes formed with the biotinylated target were captured with 25 µL streptavidin coated magnetic beads M270 (Thermo Fisher Scientific) for 15 min at 4 °C. The duration and the number of the washing steps were increased during the three rounds of selection (Table S3). The conditions of the RT-PCR, used to obtain DNA at the end of the first round were as follows: an initial denaturation step at 98°C for 30 s, followed by 35 cycles of 10 s at 98°C, 30 s at 61°C, and 10 s at 72°C with a final elongation step of 5 min at 72°C. For the following rounds, the program was the same, with 25 cycles for round 2 and 25 cycles for rounds 3 and 4.

### 2.3. Next Generation Sequencing

Each pool of DNA sequences obtained after selection were amplified and purified, adapters were added and amplicons were sequenced at the IRIC’s Genomics Core Facility at Montreal on a MiSeq 300 cycles Micro v2 (PE150). The reads obtained were assembled and analyzed to cluster sequences using a software pipeline developed previously for antibodies [29]. For practicality, we arbitrarily chose 34 Affitin sequences (AF1 to AF34), representative of the diversity of clusters, for protein production and *in vitro* characterization.

### 2.4. Enzyme-linked immunosorbent assay (ELISA)

An ELISA was designed for evaluating the specificity of selected binders for RBD. A Maxisorp plate (Nunc) was coated with 100 µL of 66 nM neutravidin and blocked with 300 µL of PBS containing 0.5 % BSA, followed by the addition of 100 µL of the biotinylated RBD at 150 nM. Then, 100 µL of 50 and 500 nM Affitins were added to the wells. Affitins were also tested for non-specific binding against directly coated BSA, streptavidin and neutravidin (66 nM). Bound Affitins were detected with 100 µL anti-MRGS-His6 antibody (Qiagen, dilution 1:5000) conjugated with horseradish peroxidase, using 100 µL *ortho*-phenylenediamine (1 mg.mL^-1^ OPD, 0.05 % H_2_O_2_, 100 mM sodium citrate pH 5.0) as substrate. The absorbance at 450 nm was recorded with a plate reader (Tecan infinite M200 Pro). All steps were performed at room temperature with 1 h of incubation in 100 µL PBS pH 7.4 before the addition of Affitins or PBS pH 7.4 containing 0.1 % Tween 20 for the following steps. Unbound molecules were removed by washing with the corresponding buffer.

### 2.5. Affinity measurements by BioLayer Interferometry (BLI) and Surface Plasmon Resonance (SPR)

An OctetRed384 BLI instrument (Sartorius) was used to determine affinities of monomeric, dimeric and trimeric Affitins for RBD. The biotinylated RBD was immobilized on streptavidin sensors and PBS containing 0.005% Tween-20 was used as interaction buffer. Data were analyzed using the Octet Analysis Studio Software. The affinities of the Affitin-Fc fusions were determined with a Biacore T200 instrument (GE Healthcare). Sensor chips SA, sensor chips CM5 and HBS-EP (0.01 M HEPES, pH 7.4, 0.15 M NaCl, 0.005% (v/v) surfactant P20, 3 mM EDTA) running buffer were purchased from GE Healthcare. Single Cycle Kinetics mode (SCK) was used for all SPR experiments. Biotinylated RBD were captured on SA chip by a capture strategy according to the supplier’s recommendations. The Affitin-Fc fusions were diluted in HBS-EP buffer at concentrations ranging from 0.62 to 50 nM and injected over the RBD-coated chip. ACE2 protein were covalently coupled to CM5 chip by amine coupling strategy according to the supplier’s recommendations. The RBD and spike Proteins were diluted in HBS-EP buffer at concentrations ranging from 0 to 500 nM and 0 to 1000 nM, respectively and injected on the ACE 2-Fc coated chip. Flow rate was set up at 30 µL/min and association and dissociation were allowed for 2 and 10min, respectively. Rmax value (RU), *k* _on_ (M^−1^·s^−1^), *k* _off_ (s^−1^), and *K*_D_ (M) were calculated from kinetic sensorgrams using the Langmuir model adapted for SCK analysis with the Biacore T200 Biaevaluation Software 3.1.

### 2.6. Nano differential scanning fluorimetry (NanoDSF)

#### 2.6.1. Thermal stability

Ten microliters of samples at 113 µg/mL were loaded into NanoDSF high sensitivity grade capillaries (NanoTemper Technologies, Munich, Germany) and installed on capillary array on the Prometheus NT.48 NanoDSF instrument (NanoTemper Technologies, Munich, Germany). The temperature was increased from 20 °C to 95 °C with a linear thermal ramp at a 1 °C/min rate. Changes in fluorescence of tryptophan at 330 and 350 nm due to denaturation of the proteins were recorded at 10 datapoints per minute rate. The 350/330 nm ratio of the two wavelengths were plotted against temperature and the first derivative analysis allowed the determination of Tm using the PR. ThermControl Software (NanoTemper Technologies, Munich, Germany). The Prometheus NT.48 also monitors the aggregation of the samples during heating.

#### 2.6.2. Stability to repeated freeze/thaw cycles

Samples were subjected to several freeze-thaw cycles of 1 hour at -80°C before they were analyzed on a Prometheus NT.48 NanoDSF instrument (NanoTemper Technologies, Munich, Germany) as described above.

#### 2.6.3. Mass photometry analysis

Mass photometry measurements were performed with a Two MP instrument (Refeyn) in a well on a glass slide according to the supplier’s instructions. Filtered buffers, samples and calibration standards were equilibrated at room temperature before use. Focus of the objective was done with 10 µL of reference buffer PBS pH7.4. Ten µL of sample at a concentration of 20 nM were added to the drop, mixed by pipetting up and down 3-4 times, to obtain a final concentration of 10 nM. The acquisition software AcquireMP recorded data for 1 minute. Analysis was performed using DiscoverMP software. Calibration was done with MFP1 proteins standards (Refeyn, MP-CON-41033) and with BSA protein (Sigma, A2153).

### 2.7. Cells

U2OS (Cat# HTB-96) cells were obtained from ATCC and transduced to express ACE2. Cells were cultured in DMEM (Dulbecco’s Modified Eagle Medium) without phenol red, supplemented with 10% of Fetal Calf Serum (FCS), 1% penicillin/streptomycin and GLUTAMAX. U2OS-ACE2 GFP1–10 or GFP 11 cells, also termed S-Fuse cells, become GFP+ when they are productively infected by SARS-CoV-2. S-Fuse cells have been described previously [30,31].

### 2.8. Pseudo particles and virus strains

Pseudotyped viruses were produced by transfection of 293T cells as previously described [32]. Briefly, cells were co-transfected with plasmids encoding for lentiviral proteins, a luciferase reporter and the SARS-CoV-2 S plasmid coding for spike proteins from the indicated variants. Pseudotyped virions were harvested at days 2-3 post-transfection. Production efficacy was assessed by measuring infectivity or the viral protein p24 concentration.

The reference D614G strain (hCoV-19/France/GE1973/2020) was supplied by the National Reference Centre for Respiratory Viruses hosted by Institut Pasteur and headed by Sylvie van der Werf. This viral strain was supplied through the European Virus Archive goes Global (EVAg) platform, a project that has received funding from the European Union’s Horizon 2020 research and innovation program under grant agreement number 653316. The variant strains were isolated from nasal swabs on Vero cells and amplified by one or two passages on Vero cells. The B.1.351 strain (CNR 202100078) originated from an individual in Créteil (France) who provided informed consent for the use of his biological materials. The P.1. strain (TY7-501), first identified in Brazil, was obtained from Global Health security action group Laboratory Network. The Omicron strain BA.1 was supplied and sequenced by the NRC UZ/KU Leuven (Belgium). The EG.5.1.3 strain (hCoV-19/France/BRE-IPP15906/2023) was supplied by the National Reference Centre for Respiratory Viruses hosted by Institut Pasteur (Paris, France) The human sample was provided by Dr F. Kerdavid from Laboratoire Alliance Anabio, Melesse (France). The JN.1 strain (hCoV-19/France/HDF-IPP21391/2023) was supplied by the National Reference Centre for Respiratory Viruses hosted by Institut Pasteur. The human sample was provided by Dr Bruno Foucault from Laboratoire Synlab Normandie Maine, La Ferté Macé (France). Individuals provided informed consent for the use of their biological materials.

Titration of viral stocks was performed on Vero E6 cells, with a limiting dilution technique allowing a calculation of the 50% tissue culture infectious dose, or on S-Fuse cells. Viruses were sequenced directly on nasal swabs and after one or two passages on Vero cells.

### 2.9. Neutralization tests

Affitins were prepared at varying concentrations through serial 1:10 dilutions in a 96-well white-edged plate. Each well contained 25 µL of diluted Affitins, and all conditions were tested in duplicate. Lentiviral particles containing the luciferase cassette and pseudotyped with the SARS-CoV-2 spike protein were diluted to the desired concentration (60ng/ml) and 25 µL of the viral suspension was added to each well containing 25 µL of pre-diluted Affitins. The virus-Affitin mixture was incubated for 30 minutes at room temperature to allow potential interaction. After the incubation period, 50 µL of the U2-OS cell suspension (10,000 cells in DMEM) was added to each well, bringing the total volume to 100 µL per well.

Seventy-two hours post-infection, viral entry was assessed using the ONE-Glo™ Luciferase Assay System (Promega). To each well, 50 µL of luciferase substrate was added. The plate was incubated for 5 minutes in the dark to ensure complete reaction, and luminescence was measured using a Victor plate reader. The percentage of neutralization was calculated using the luminescence intensity with the following formula: 100 × (1 − (value with Affitin − value in ‘non-infected’)/(value in ‘no Affitin’ − value in ‘non-infected’)). Neutralizing activity of each Affitin was expressed as the Half maximal inhibitory concentration (IC_50_). IC_50_ values were calculated with a reconstructed curve with the Prism software (Nonlinear regression model: log(inhibitor) vs. response-variable slope-four parameters with top constrained to 100% and bottom constrained to 0%) using the percentage of neutralization at each concentration.

### 2.10. S-Fuse neutralization assay (Pasteur)

U2OS-ACE2 GFP1–10 or GFP 11 cells, also termed S-Fuse cells, become GFP+ when they are productively infected by SARS-CoV-2. Cells were mixed (ratio 1:1) and plated at 8 × 10^3^ per well in a μClear 96-well plate (Greiner Bio-One). The indicated SARS-CoV-2 strains were incubated with serially diluted monoclonal antibodies or Affitins for 15 min at room temperature and added to S-Fuse cells. Eighteen hours later, cells were fixed with 2% PFA (Electron microscopy cat# 15714-S), washed and stained with Hoechst (dilution of 1:1000, Invitrogen cat# H3570). Images were acquired using an Opera Phenix high-content confocal microscope (PerkinElmer). The GFP area and the number of nuclei were quantified using the Harmony software (PerkinElmer). The percentage of neutralization was calculated using the number of syncytia as value with the following formula: 100 × (1 − (value with Affitin− value in ‘non-infected’)/(value in ‘no Affitin’ − value in ‘non-infected’)). Neutralizing activity of each Affitin was expressed as the Half maximal inhibitory concentration (IC50). IC50 values were calculated with a reconstructed curve using the percentage of neutralization at each concentration.

### 2.11. Structure prediction

Interaction models of SARS-CoV-2 spike beta with AF11-11-11 and AF24-24-24 were created with Alphafold3 server [33]. For the two models, each sequence of SARS-CoV-2 spike B1.351 (Table S1) trimer and AF11-11-11 and AF24-24-24 (Table S1) were input as separate protein entities. Each obtained complex was then optimized in MOE (*Molecular Operating Environment (MOE)*, 2024.0601 Chemical Computing Group ULC, 910-1010 Sherbrooke St. W., Montreal, QC H3A 2R7, **2025**) using the Protein Preparation tool, and hydrogens added with Protonate 3D using standard parameters. The complexes were then minimized in MOE using the Amber10:EHT forcefield in implicit water with a dielectric constant of 80 and an RMS gradient of 0.0001 kcal/molÅ^2^. The structures were then visually inspected and analyzed using the protein contact tool. Figures were also created with MOE.

## 3. Results

To harness the combined advantages of Affitins, which are small, stable and easily modifiable proteins, and their generation entirely in vitro by ribosome display, we have setup a pipeline for the engineering of multimeric Affitins with potent neutralization properties (Figure 1). As a case study, we choose to neutralize SARS-CoV-2 a widely studied virus. This pipeline begins with the generation of two highly diverse Affitin libraries specifically designed to recognize targets through distinct binding modes [25,28]. Thus, potential interactions with the target can occur either through a randomized flat surface (library L5) or through a combination of a flat surface and a small, artificially extended flexible loop (library L6), enabling the identification of binders suited to various epitope geometries. Next, ribosome display is used to drive selections against the target of interest, here RBD of the spike protein from SARS-CoV-2. The DNA outputs of selections are then sequenced by NGS and analyzed to cluster them by families in order to study also weakly represented selected sequences. Representative sequences of clusters are then obtained by gene synthesis, allowing their expression and characterizations, such as neutralization properties. Next, the most promising candidates are assembled via GoldenGate approach to generate various combinations of dimers and trimers. Finally, depending on the properties sought, another level of multimerization is reached via fusion of the best trimers candidates to a human Fc fragment, thereby resulting in Affitins hexamers.

**Figure 1.**
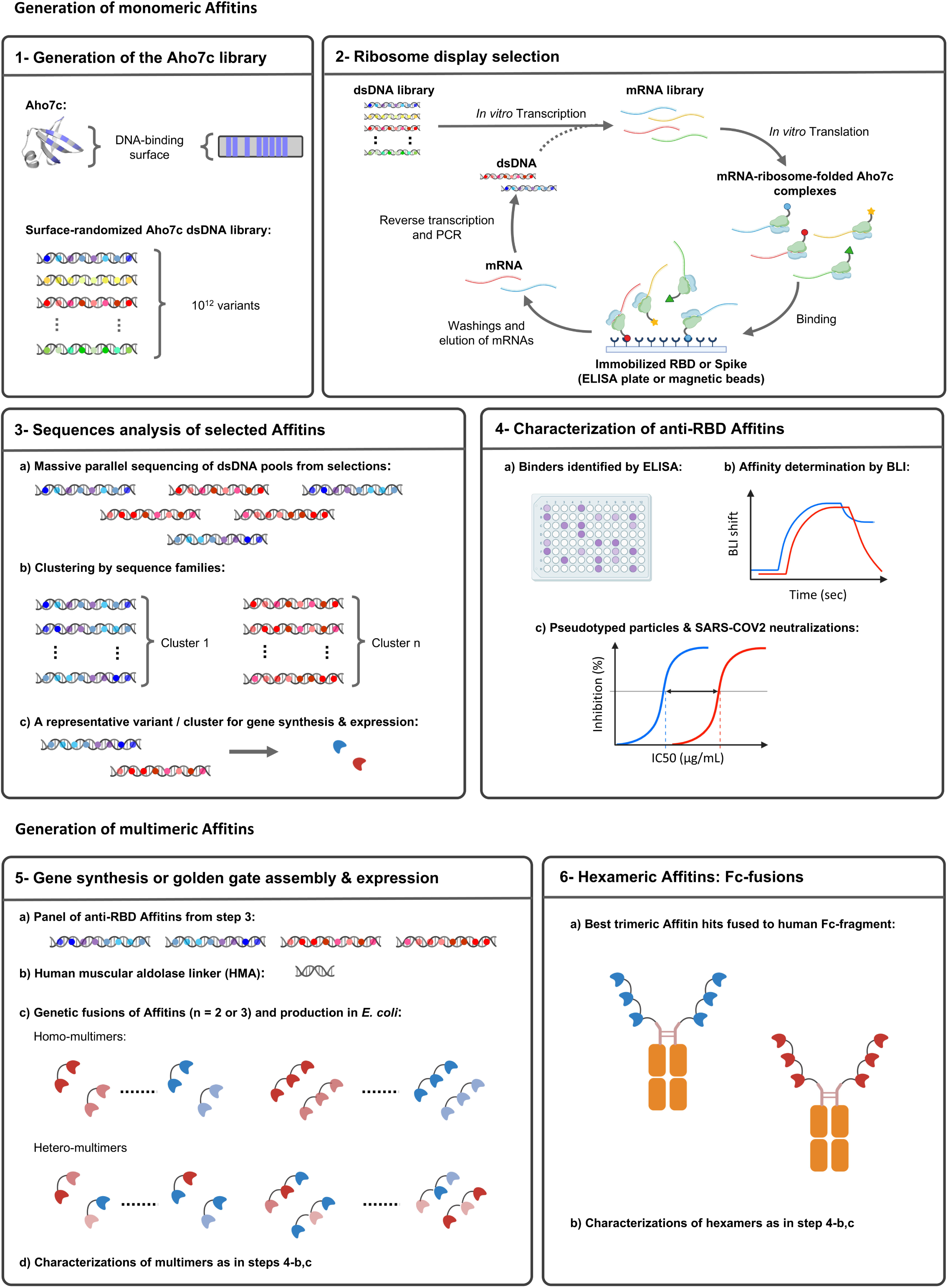
Workflow of the pipeline for generating neutralizing multivalent Affitins. (**1**) Large synthetic DNA libraries are created in vitro by randomizing specific positions within the Aho7c sequence (depicted in blue on the structure and gene representation). This is achieved using degenerate oligonucleotides encoding NNS and NHK codons. (**2**) These DNA libraries are subjected to in vitro selection against the target of interest, here spike and RBD, using ribosome display. (**3**) Post-selection, the pools of potential binders are analyzed through Next-Generation Sequencing (NGS). The sequences are clustered to identify representative clones corresponding to the sequence families in the selection outputs. (**4**) The representative Affitins are evaluated for their binding properties using ELISA and Biolayer Interferometry (BLI), as well as their neutralization potency. (**5**) Lead Affitins are assembled into homo- and heterodimers and trimers using GoldenGate assembly with the Human Muscular Aldolase (HMA) linker. (**6**) Trimers are further fused to a human Fc fragment to generate higher-order multimers. All multimers are thoroughly characterized for their binding affinity and neutralization properties as multivalent constructs. Created in https://BioRender.com.

### 3.1. Selection of anti-RBD Affitins

As the infection of host cells by SARS-CoV-2 requires interaction of the RBD region of its spike protein with the membrane receptor ACE2, we aimed to isolate Affitins against the RBD domain. The first two rounds of selection by ribosome display were performed with the spike protein as bait. The third round was done using RBD as target to select Affitins that only bind to this region. A polyclonal ELISA test performed on in vitro translated pools of selected clones from both L5 and L6 libraries was positive already from round 3 (Figure S2-A). After the 3rd round of selection, we found that 69% and 94% of tested clones from library L5 and L6, respectively, were positive in ELISA for specific binding to RBD domain (i.e. with a specific/aspecific signal ratio per well higher than 10).

The pools of selections were then submitted to massive parallel sequencing (NGS) and the sequences obtained were analyzed to cluster them. The Affitins representative of these clusters were produced for their characterization. The ELISA test results in Figure S2-B demonstrate that 9 Affitins were able to specifically recognize the RBD at concentrations of 500 nM as evidenced by the absence of binding to NeutrAvidin used to immobilize RBD. We identified 8 Affitins isolated from library L5 (AF3, AF4, AF5, AF6, AF7, AF8, AF11, and AF15), and 1 from L6 library (AF24), with quite different sequences (Figure S3). The Affitins AF4, AF5, AF7, AF8, AF11, AF15, and AF24 showed a specific binding to RBD (no binding to BSA and NeutrAvidin) and an OD450 signal of at least 1 when tested at a concentration of 50 nM, and were selected for further characterization.

### 3.2. Characterization of monomeric Affitins

The Affitins were efficiently produced, with purification yields ranging from 16 mg/L to 119 mg/L of *E. coli* culture, averaging 53 mg/L. All Affitins eluted from the size-exclusion chromatography column as a monomer peak and migrated according to their molecular weights on a SDS-PAGE gel (Figure S4).

We then determined their affinity for the recombinant RBD by BLI (Figure 2A, Table S4). All Affitins exhibited affinities within the nanomolar range, with AF4 demonstrating the highest affinity at 15.6 nM, while AF24 showed the lowest affinity at 603 nM.

**Figure 2.**
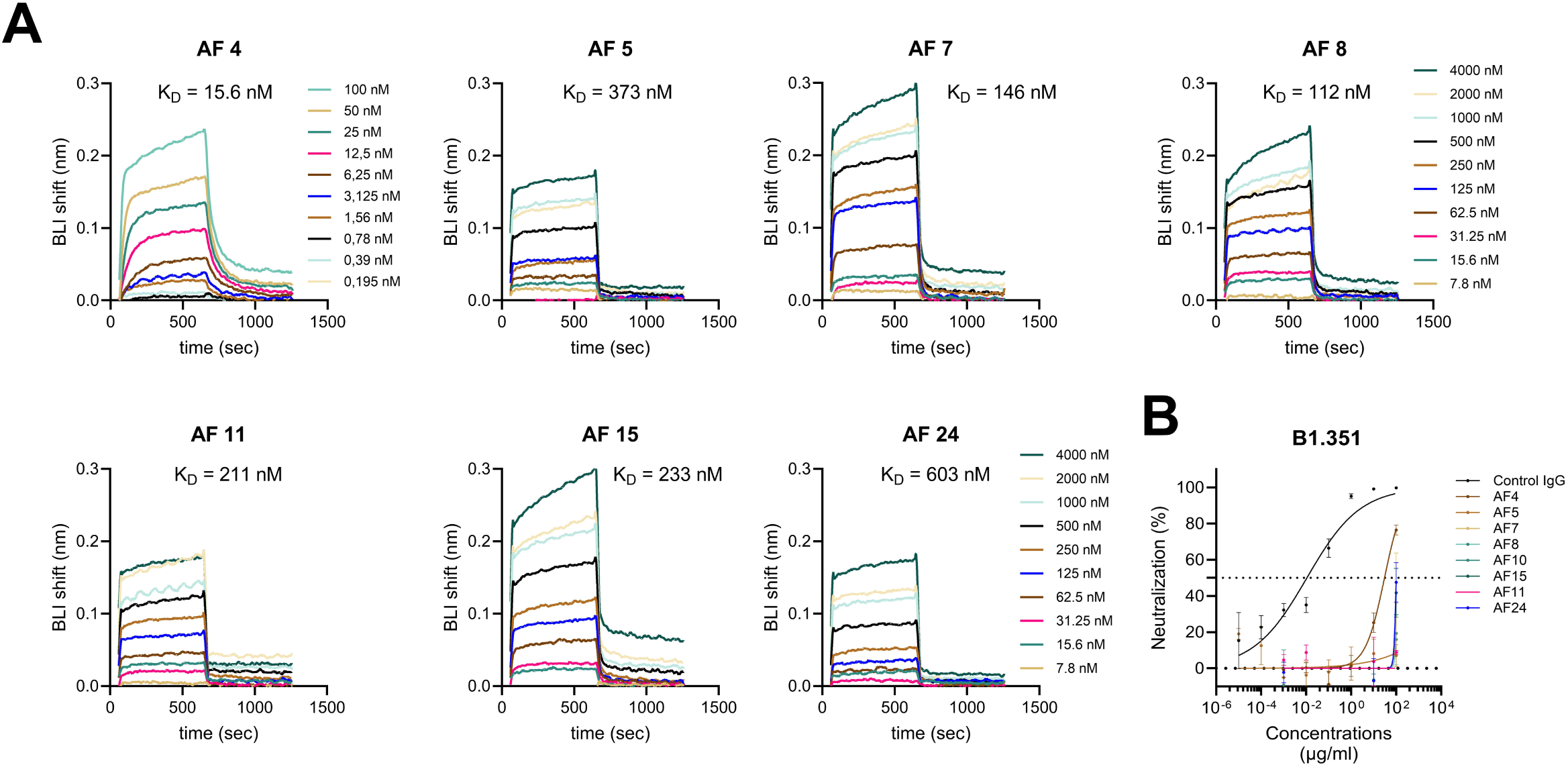
Characterization of the monomeric Affitins. (**A**) Study by BLI of the binding kinetics of Affitins at various concentrations to RBD immobilized on streptavidin-sensor. (**B**) B1.351 pseudovirus neutralization by Affitins (n = 3).

An analysis by NanoDSF showed that Affitins AF4, AF5, AF11 and AF15 were extremely stable, with no thermal denaturation observable up to 95°C (Table S5). A denaturation was however observed for Affitins AF7, AF8, and AF24, with melting points (Tm) of 64.52°C, 64.66°C, and 84.68°C, respectively. Notably, no aggregation was observed for any of the Affitins across the temperature range of 20°C to 95°C at the tested concentration of 113 µg/mL. In contrast, the mAb Cv2.1169 used in most of our neutralization studies displayed two thermal denaturation transitions, with a first event at 70.23°C, and a second one at 83.03°C. An aggregation process occurred around 85.22°C after the second denaturation event (Table S5).

Next, we tested the neutralization properties of monomeric Affitins against B1.351 pseudovirus from which RBD was used for selections (Figure 2B). The Affitin AF4 demonstrated the highest ability to neutralize cellular infections, achieving an effective concentration (IC_50_) of approximately 50 µg/mL, equivalent to about 6 µM. In this experiment, the control mAb was far more efficient with an IC_50_ of 10^-2^ µg/mL (about 67 pM).

### 3.3. Characterization of dimeric and trimeric Affitins

Since the monomeric form of the Affitins showed weak or no neutralization effect on B1.351 pseudovirus, even for AF4 which has the highest affinity for RBD, we sought to investigate whether multimerization could enhance their effectiveness. To this end, we produced homodimeric forms of AF4, AF5, and AF24, representing high, medium, and low affinity anti-RBD Affitins, respectively. We also produced the heterodimer AF5-15 (Figure S4), which combines Affitins with moderate affinities that showed no neutralization effect in their monomeric forms. The data obtained for the neutralization of B1.351 pseudovirus shown in Figure 3A indicate that homodimerization improved significantly IC_50_. For instance, for dimeric AF5 and AF24 we could determine IC_50_ of 93 and 4.5 µg/mL (5.6 and 0.27 µM), respectively, while monomeric AF5 and AF24 did not shown a neutralization effect (Figure 2B). Surprisingly, AF4 which was the only Affitin able to neutralize as a monomer (IC_50_ = 50 µg/mL, 6.0 µM), did not show the best neutralization effect among the dimers tested (IC_50_ = 15,6 µg/mL, 0.92 µM). Unexpectedly, the heterodimer AF5-15 demonstrated nearly equivalent potency to the most effective dimer, AF24-24 (IC_50_ of 4.5 µg/mL, 0.27 µM), achieving an IC_50_ of 5.9 µg/mL, 0.36 µM. We determined that the dissociation constant of AF5-15 for RBD was 9.5 nM, representing a 39-fold and 25-fold improvement in binding affinity compared to monomeric AF5 and AF15, respectively, mostly via a decrease of the dissociation rate (Figure S5, Table S6).

**Figure 3.**
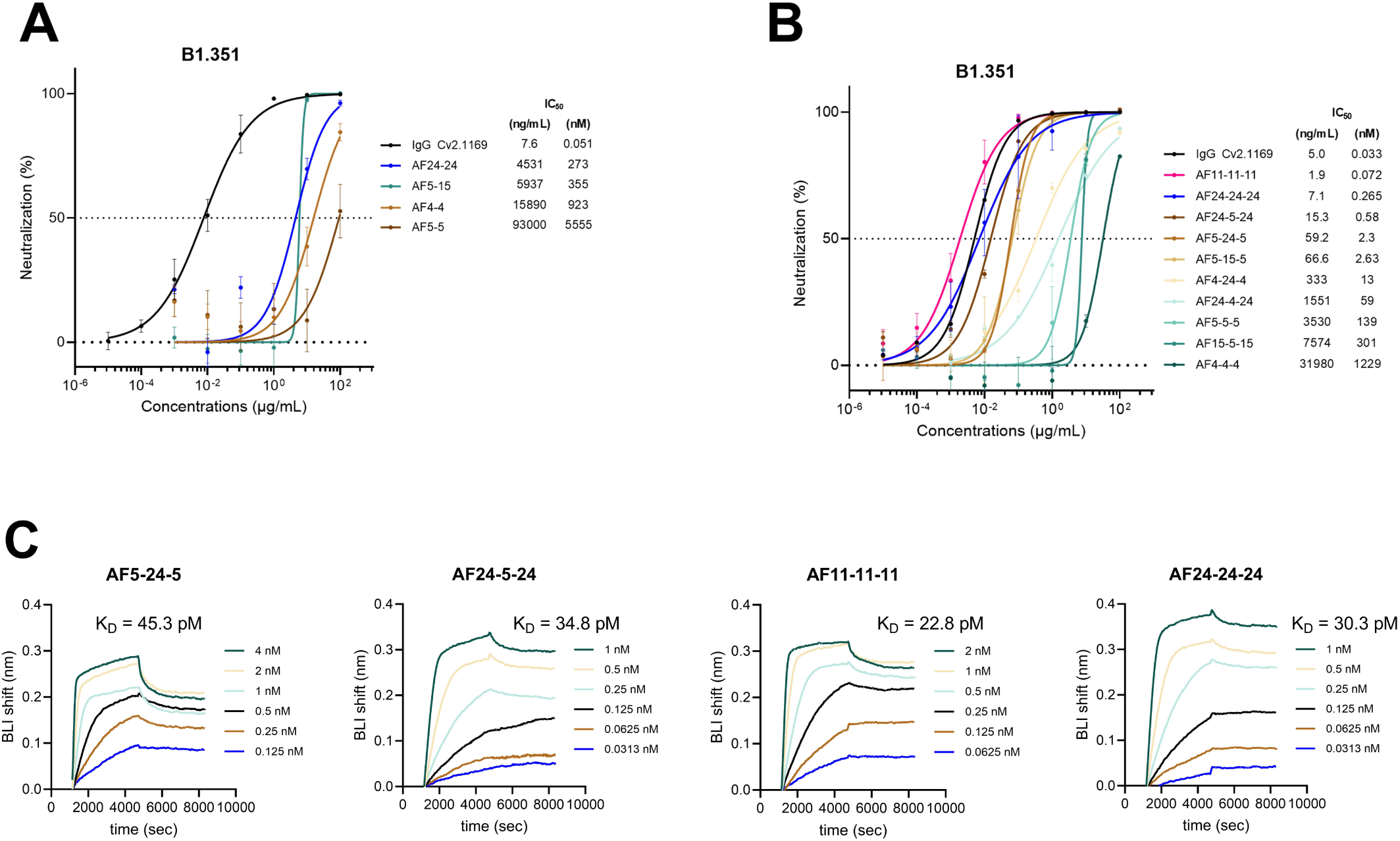
Characterization of dimeric and trimeric Affitins. Neutralizations of pseudovirus B1.351 by (**A**) dimeric (n=2) and (**B**) trimeric Affitins (n = 3). The monoclonal antibody Cv2.1169 was used as positive control for neutralization [34]. (**C**) BLI of the binding kinetics of Affitins RBD immobilized on streptavidin-sensor.

Overall, these data show that dimerization is efficient to significantly improve the neutralization activity of Affitins, probably via an avidity effect as observed for AF5-15. However, as the control antibody Cv2.1169 still showed a higher neutralization activity than dimers with an IC_50_ of 7.6 ng/mL, 51 pM (Figure 3A), the dimers were not further characterized. Given that the spike protein is a homotrimer, we investigated the properties of trimeric anti-RBD Affitin constructs. We focused on the most stable Affitins from the L5 library, as identified through NanoDSF analysis: AF4, AF5, AF11, and AF15 (Tm > 95°C; Table S5), excluding the Affitins AF7 and AF8. Additionally, AF24 was included for characterization as the sole representative of the L6 library which has the potential to exhibit a distinct binding mode to the target, thanks to the artificially extended loop engineered in the L6 library [13]. Accordingly, a set of ten representative trimers were produced using Affitins AF4, AF5, AF11, AF15, and AF24 (Figure S4).

As shown in Figure 3B, trimerization had a pronounced impact on neutralization activity for B1.351 pseudovirus, particularly for AF11-11-11 and AF24-24-24, which achieved IC_50_ values of 1.9 ng/mL (72 pM) and 7.1 ng/mL (265 pM), respectively. The neutralization activity of AF11-11-11, closely matches that of the control antibody Cv2.1169 (IC_50_ = 5.0 ng/mL, 33 pM). Interestingly, AF5-5-5 demonstrated a 40-fold improvement in neutralization activity compared to AF5-5, while AF24-24-24 exhibited an impressive 1030-fold enhancement against B1.351 pseudovirus compared to AF24-24. Similarly, extending AF5-15 by adding AF5 to its C-terminus (resulting in AF5-15-5) led to a 135-fold improvement. However, adding AF15 to its N-terminus (forming AF15-5-15) showed no significant effect (1.2-fold).

As observed for dimerization, trimerization significantly improved binding affinities, with all tested trimers achieving dissociation constants within the picomolar range (Figure 3C, Table S6). Among the tested constructs, AF11-11-11 and AF24-24-24 demonstrated the strongest binding affinities, with dissociation constants estimated at 22.8 pM and 30.3 pM, respectively. This increase in affinity was primarily driven by a 3- to 4-order-of-magnitude reduction in dissociation rates compared to the monomeric forms. These findings further reinforce the correlation between enhanced binding affinity and neutralization potency, as these two trimers also exhibited the highest efficacy in neutralization assays.

We then evaluated the cross-neutralization activities of the four most potent trimers against different pseudovirus (Figure 4A). As expected, all were able to neutralize B1.351 that hosts the RBD used for selections of Affitins. The neutralizing activity was dependent on the trimer/pseudovirus tested. For instance, AF24-24-24 displayed cross-neutralization for all pseudovirus variants tested in the nanomolar range (from IC_50_ = 1.45 nM for B1.617.2 to IC_50_ = 107 nM for BA.1), but for higher concentration than with the control antibody Cv2.1169. AF24-5-24 was able to neutralize all but BA.1 pseudovirus (when measurable, with IC_50_ = 1.57 nM for D614G to IC_50_ = 23 nM for B1.1.7), the genetically most distant one from B1.351 in our test. AF5-24-5 had a profile similar to AF24-5-24. AF11-11-11 was the less cross-reactive trimer, since it could not neutralize D164G, B1.1.7, B1.617.2, and BA1, demonstrating a high specificity.

**Figure 4.**
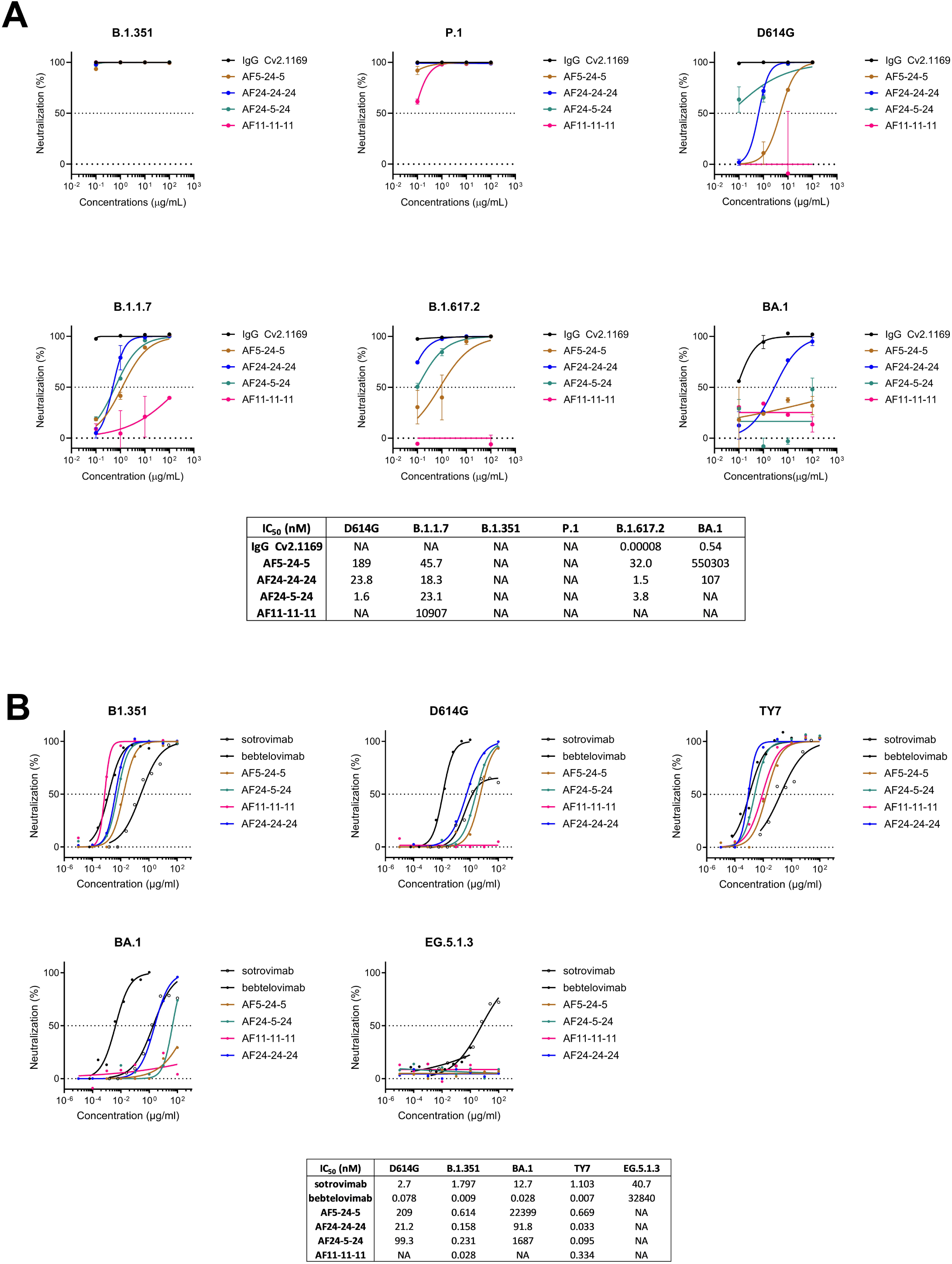
Neutralization properties of trimeric Affitins. **(A)** Neutralizations of pseudovirus B1.351, P.1, D614G, B1.1.17, B1.617.2 and BA.1 by trimeric Affitins (n = 2). The monoclonal antibody Cv2.1169 was used as positive control for neutralization [34]. (**B**) Neutralizations of SARS-Cov-2 virus B1.351, D614G, TY7, BA.1 and EG5.1.3 (n = 1). Survival of human cells in absence of virus used for the test is represented as a green line. sotrovimab [47] and bebtelovimab [48] were used as positive controls for neutralization.

Next, we evaluated the neutralization effects of the trimers against various virus strains of SARS-CoV-2 (Figure 4B). AF11-11-11 emerged as the most potent trimer against the B1.351 strain (IC_50_ = 28 pM), comparing well with the control antibody bebtelovimab (IC_50_ = 9 pM), and outperforming sotrovimab (IC50 = 1800 pM) . However, its activity dropped significantly when tested against other virus strains, with the exception of the TY7 strain (IC_50_ = 334 pM). In contrast, AF24-24-24 demonstrated the highest potency and cross-reactivity among the trimers with IC_50_ from 33 pM for TY7 to IC_50_ = 92 nM for BA.1, although it failed to neutralize the EG5.1.3 strain. AF5-24-5 and AF24-5-24 exhibited a neutralization profile similar to that of AF24-24-24 but were less potent in their overall activity. Interestingly, none of the trimers exhibited toxicity toward the human cells used in the neutralization assays across the tested concentration range: up to 100 µg/mL, about 4 µM (Figure S6).

Since AF11-11-11 showed the highest specific neutralization activity against the B1.351 strain and AF24-24-24 proved to be the most potent and cross-reactive trimer, both were further analyzed. The competitive binding assays presented in Figure S7 demonstrated that the binding of AF11-11-11 almost entirely inhibited the binding of AF24-24-24, and this effect was reciprocal. This result suggests that both binders target the same epitope or recognize overlapping epitopes. Thermal stability measurements by NanoDSF, as presented in Table S7, revealed a unique thermal transition with associated Tm values of 75.3°C and 84.01°C for AF11-11-11 and AF24-24-24, respectively, indicating that the trimers are thermally stable. However, these findings suggest that multimerization can impact stability, as the monomeric forms of AF11 did not exhibit thermal denaturation even at temperatures exceeding 95°C. In contrast, the monomeric and trimeric forms of AF24 exhibited comparable thermal stability, with no significant difference observed between the two formats. Interestingly, no aggregation was observed for any of the trimers across the temperature range of 20°C to 95°C. Furthermore, repeated freeze/thaw cycles did not alter the Tm values, and no aggregation was detected, demonstrating the robustness of these trimers. Overall, these results further supported our trimerization strategy.

### 3.4. Characterization of Affitin-hFc1 fusions

Given that dimerization and trimerization significantly and gradually enhanced neutralization potency, we explored whether an even greater effect could be achieved through a higher degree of multimerization. To this aim we fused either AF11-11-11 or AF24-24-24 trimer to a human Fc1 fragment (noted hFc1) to generate dimers of trimers, and thus homohexameric Affitins: (AF11-11-11)_2_-hFc1 and (AF24-24-24)_2_-hFc1. These fusions were produced in Expi293 with high yield of production, 69 and 106 mg/L of culture, respectively. The SDS-PAGE analysis showed that the assembly of the Fc region occurred via disulfide bond formation (Figure S4C). The molecular weights of purified (AF11-11-11)_2_-hFc1 and (AF24-24-24)_2_-hFc1, as determined by mass photometry (106 kDa) were in agreement with the theoretical values of 101 and 102 kDa without glycosylation, respectively (Figure S4E). Furthermore, according to mass photometry both proteins were monodisperse supporting a high degree of purity. We analyzed their affinity for immobilized RBD using SPR (Figure 5A, Table S8), determining dissociation constants of 23.4 pM and 13.7 pM, respectively. Unlike the significant affinity gains observed with monomer-to-dimer or dimer-to-trimer multimerizations, according to SPR measurements the increase in affinity was negligible for (AF11-11-11)_2_-hFc1 and modest for (AF24-24-24)_2_-hFc1, with only a 2.2-fold improvement.

**Figure 5.**
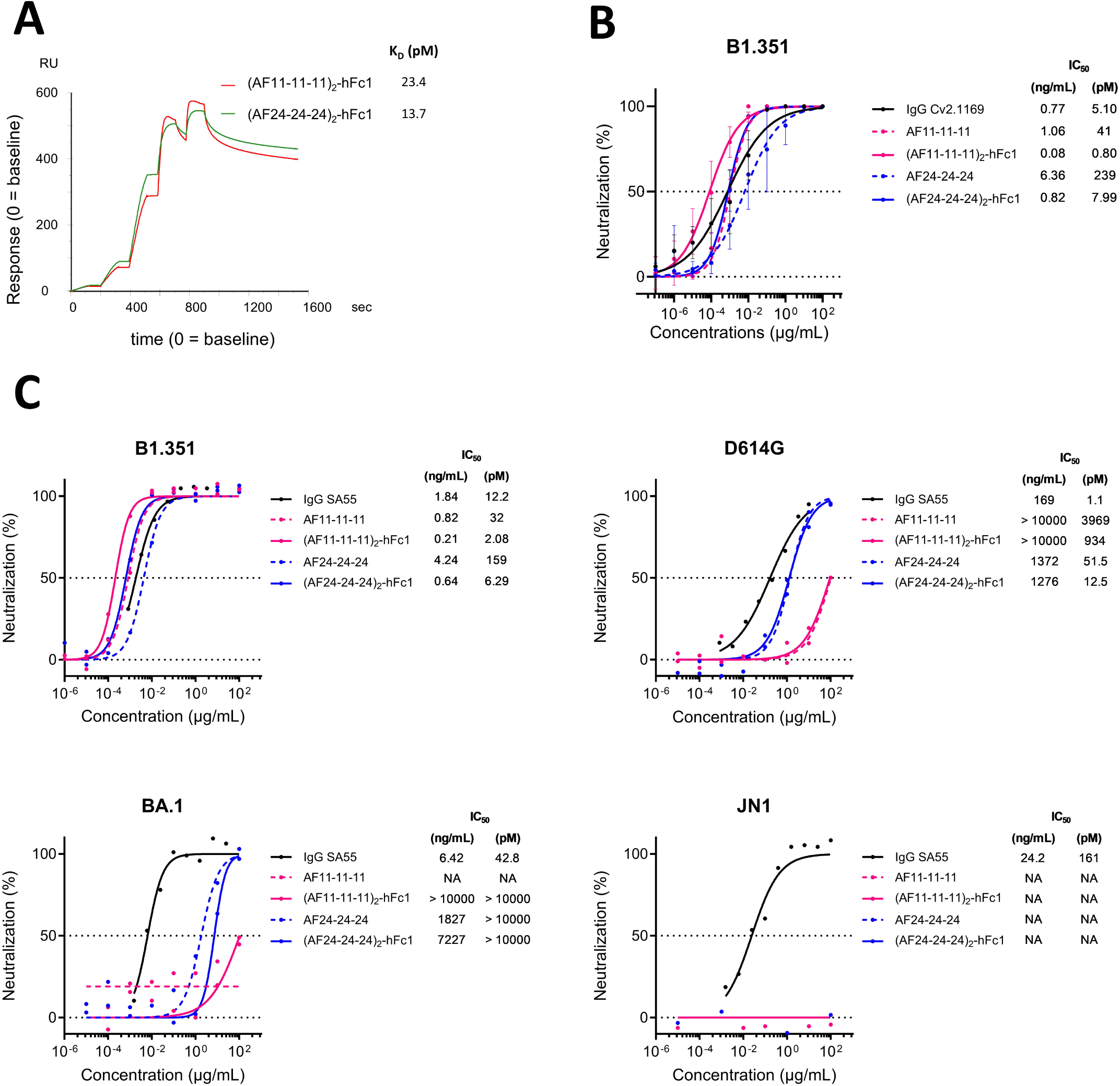
Characterization of hexameric Affitins. **(A)** SPR of the binding kinetics of Affitins to RBD immobilized on streptavidin-sensor. The analysis was performed by the single-cycle kinetics (SCK) method with no regeneration of the sensor between injections of Affitins. (**B**) Comparison of neutralization activities of trimeric and hexameric Affitins for pseudovirus B1.351 (n = 4). The monoclonal antibody Cv2.1169 was used as positive control for neutralization. (**C**) Same as in (**B**) but for SARS-Cov-2 virus B1.351, D614G, BA.1 and JN1. Monoclonal antibody SA55 was used as positive control for neutralization [35].

We then examined whether this relatively modest increase in affinity would impede any potential improvement in neutralization potency. The Figure 5B shows neutralization results obtained for trimers, hexamers and the control antibody Cv2.1169 [34] against pseudovirus B1.351. The IC_50_ determination revealed that (AF11-11-11)_2_-hFc1 was 51 times more potent than AF11-11-11 (0.8 vs 41 pM), while (AF24-24-24)_2_-hFc1 demonstrated a 30-fold increase in potency compared to AF24-24-24 (8.0 vs 239 pM). Interestingly, (AF11-11-11)_2_-hFc1 outperformed the control antibody, exhibiting a 6-fold greater potency (0.8 vs 5.1 pM).

Next, we studied how behaved the trimers and hexamers to neutralize viruses (Figure 5C). The results obtained with B1.351 SARS-CoV-2 matched closely those obtained for B1.351 pseudovirus. The hexamer fusions exhibited enhanced potency against B1.351 strain compared to their corresponding trimers, with a 15-fold increase for AF11 multimers (2.1 vs 32 pM) and a 25-fold increase for AF24 multimers (6.3 vs 159 pM). Remarkably, (AF11-11-11)_2_-hFc1 exhibited a 6-fold higher neutralization activity compared to the IgG SA55 [35], further supporting the effectiveness of our multimerization approach for Affitins. Both constructs were capable of at least partially neutralizing several SARS-CoV-2 strains, including B1.351, D614G, and BA.1. However, neither construct showed any neutralization activity against the JN1 strain, the most genetically distant strain from B1.351 in our tests. As for trimers, no toxicity could be observed with hexameric Affitins up to a concentration of 100 µg/mL, 1 µM (Figure S6).

These hexameric Affitins were also confirmed to be thermostable, as determined by NanoDSF (Table S7). The AF11-based hexamer exhibited a Tm value of 68.78°C, lower than that of AF11-11-11 (75.3°C). The AF24-based hexamer had a Tm value of 68.74°C, which was lower than the Tm of AF24-24-24 (84.01°C). For these Fc-fusion constructs, a second thermal transition was observed, with Tm values of 81.67°C and 82.51°C for (AF11-11-11)_2_-hFc1 and (AF24-24-24)_2_-hFc1, respectively, likely corresponding to the denaturation of domain 3 of the Fc region. Notably, repeated freeze/thaw cycles did not alter the Tm values, and no aggregation was detected, demonstrating the robustness of these Fc-fusion constructs.

### 3.5. Modeling the interaction between Affitins and the spike protein

To gain deeper insight into the possible interactions by which Affitins neutralize viruses, three-dimensional structural modeling of the AF11 and AF24 trimers in complex with the spike protein was performed. Reliable complexes were obtained for each trimer with an interface predicted template modeling (ipTM) 0.78 and predicted template modeling (pTM) 0.80 for the AF11 complex, and 0.78 ipTM and 0.79 pTM for AF24, models are presented in Figure 6. According to these models, both AF11 and AF24 bind to the spike protein in the RBD region via their amino-acids that were randomized in the libraries, and the two Affitins recognize different and distant epitopes (Figure 6). The analysis of the observed interactions between each AF11 or AF24 monomers are tendentially repeated to each spike RBD monomer (Table S9, Figure 6 & S8), therefore we focused the analysis on the interaction pattern of the randomized residues for the first Affitin monomer.

**Figure 6.**
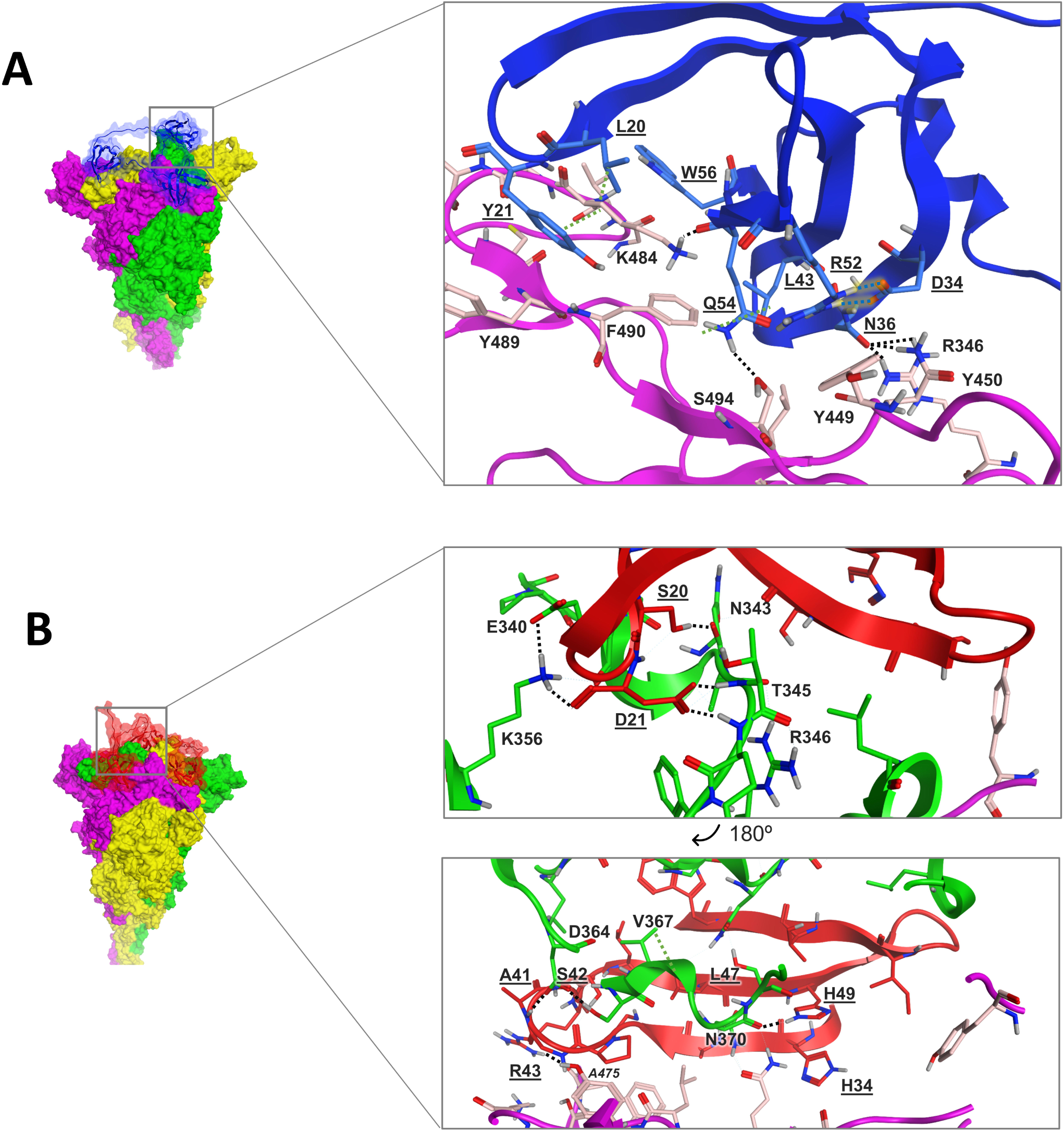
Modeling of the complexes and interactions between trimeric Affitins and the spike protein: (**A**) for AF11-11-11 and (**B**) for AF24-24-24; underlined residues - Affitins, bold residues - spike chain A and residues in italic - spike chain B; green dotted lines - hydrophobic interactions; blue dotted line with yellow glow - ionic interactions; black dotted lines - hydrogen bonds.

AF11 randomized residues (Figure S3) display key interactions that may rationalize the high affinity observed experimentally. At the beginning of the first loop in AF11 structure, L20 and Y21 have energetically favorable interactions (Table S9). The residue L20 interacts hydrophobically with the aliphatic region of K484 side chain of the RBD and Y21 with the same region and tyrosine’s aromatic ring. Other close contacts with G485, F486 and C488 also contribute favorably for the affinity of this AF11 residue. The second randomized region has residues that do not interact with spike’s RBD, like M33, and D34 which does not increase the interaction with spike due to its side chain CH_2_ group being too close to N450 polar side chain amide. However, D34 strong ionic interaction with AF11 R52 can be important to stabilize this residue and N36 positions in order to establish the strong hydrogen bond network the latter residue presents with the RBD. In fact, N36 presents a high contribution to the binding of AF11 to the RBD by hydrogen bonding with spike’s R346 and N450 side chains. L43 performs contributing hydrophobic interactions with L452 and F490 side chains, its randomized neighboring residue G45 does not present any interaction of notice. The last randomized region comprises the previously described R52, Q54 and W56. As one of the most important interactions between AF11 and spike, Q54 side chain present a hydrogen bond with S494 hydroxyl and Q54 backbone carbonyl with the side chain of K484. W56 interacts aromatically and hydrophobically with V483 side chain.

Unlike AF11-11-11 that has each Affitin monomer interacting to a single RBD, AF24 trimer sees its monomers laying in the cleft region between two RBD chains. This placement might be due to the longer loop between beta-sheets 3 and 4 making a higher surface area of contact, increasing the possibility of making stable interactions with spike. The first two randomized residues S20 and D21 are contributors for the binding of AF24. The side chain of S20 displays a hydrogen bond with N343 backbone carbonyl and D21 is one of the residues that contribute more for AF24 binding. The aspartate side chain does several charge-induced hydrogen bonds with hydroxyl group and backbone nitrogen of T345, and another with R346 backbone. Furthermore, its backbone carbonyl group also interacts with K356 side chain. On the region between H34 and H49, some of the residues are great contributors for AF24 binding particularly: A41 and S42 backbone nitrogens hydrogen bonding Asp364 side chain; R43 side chain interaction with backbone carbonyls of A475 and G476; L47 hydrophobic interaction with V367; and H49 side chain with the backbone carbonyl of N370.

Interestingly, each recognition moiety of the AF11- and AF24-trimers seems to bind to each the three RBDs from spike. These observations correlate with the large improvement both for affinities and neutralization potencies seen with trimers compared to monomers and dimers, and also the different neutralization properties of AF11-11-11 and AF24-24-24. Finally, these models validate the length of the linker used (20 amino-acids) and the interest to isolate Affitins from libraries with different designs.

## 4. Discussion

Many neutralizing IgG-type antibodies have been generated against viruses in general, with a recent example being SARS-CoV-2, which has received particular attention due to the pandemic it caused. While antibodies are potent therapeutic agents, they can present certain drawbacks due to their structural complexity and their large molecular size. Viruses frequently present repeated epitopes on their surface, making multivalent affinity molecules significantly more effective than their monovalent counterparts due to the avidity effect. A notable example is the HIV-neutralizing antibody 2G12, which, although inherently bivalent, naturally forms dimers and trimers. These multimeric forms exhibit a neutralizing potency up to 2,500 times greater than that of the monomer [36]. Similarly, artificially multivalent IgG antibodies, engineered through disulfide bridge formation or multimerizing mutations, have also demonstrated greater potency compared to their monomeric equivalents [37,38]. However, these approaches result in complex structures with very high molecular weights, which hinder their industrial-scale production and use. As a result, research into IgG multimerization to enhance therapeutic efficacy remains limited due to these bottlenecks.

Affitins provide a versatile molecular platform that is significantly simpler than IgG or VHH antibodies in terms of structure (no disulfide bonds, a single polypeptide chain), size (∼7 kDa), and stability [13,14,25]. In this work, we identified seven Affitins with affinities for the recombinant RBD spanning a broad range of K_D_ values, from 15.6 nM for AF4 to 603 nM for AF24 (Figure 2A and Figure S2). With the exception of AF4, these Affitins, despite binding to the RBD according to ELISA and BLI experiments, did not exhibit neutralizing activity against B.1.351 pseudovirus within the tested concentration range. These Affitins exhibited remarkable stability, with thermal stability values ranging from 64.52°C to over 95°C, and were efficiently produced in bacteria at high yields (16 to 119 mg/L). This made them particularly suited for engineering multimeric constructs.

To target the spike protein, which forms a homotrimer, we used a 20-amino-acid peptide spacer to link Affitins. The spike protein on the surface of SARS-CoV-2 predominantly displays a mixture of RBD conformations, with one RBD in the "up" state and two in the "down" state being the most frequently observed. In this prefusion conformation, the distances between the centers of the ACE2-binding interfaces on each RBD were measured to range between 36 and 59 Å, based on the 3D structure with PDB code 6VSB [19]. The chosen linker, capable of spanning a theoretical distance of approximately 70–76 Å in a fully extended conformation, was expected to provide sufficient flexibility for dimeric or trimeric Affitins to simultaneously engage two or three RBDs, respectively. The data obtained from BLI and neutralization experiments highlight the effectiveness of this approach (Figures 2 and 3). For example, trimerization enhanced the affinity of AF11 for the RBD by 9,253-fold, while AF24 exhibited an even greater affinity increase of 19,924-fold. AF4 was the only Affitin capable of neutralizing the B1.351 pseudovirus as a monomer, with an IC_50_ of about 6 µM. This allowed us to quantify the enhanced neutralization effects of its dimerization (IC_50_ = 923 nM, 6.5 fold) and trimerization (IC_50_ = 1.23 nM, 4.9 fold). Surprisingly, despite its initial capabilities, AF4 proved not to be the optimal candidate for designing a potent neutralizing protein. This was due to its limited improvement upon dimerization and the absence of any further enhancement in neutralization efficacy with trimerization. In contrast, monomeric forms of AF11 and AF24 were ineffective for neutralization, but when assembled into trimers, they became the most potent Affitins for neutralizing the B1.351 pseudovirus, with IC_50_ values of 72 pM and 265 pM, respectively. These findings underscore the importance of exploring a diversity of binders, as the affinity of monomers alone is not always the most critical criterion for selecting candidates for subsequent assembly.

Our pipeline addresses this by employing strategies to ensure sufficient diversity in Affitin sequences. This is achieved through next-generation sequencing (NGS) analysis of selection outputs and the use of two libraries with distinct designs (L5 and L6). Additionally, our approach facilitates the combinatorial assembly of these diverse monomers using the GoldenGate method, enabling the efficient generation of multimeric constructs with enhanced properties.

Finally, we achieved a higher level of multimerization by fusing the trimers to a human Fc1 fragment, thereby generating hexamers. Surprisingly, this fusion did not significantly enhance the affinities of the hexamers compared to their corresponding trimers, with no or only modest improvements of 0.97-fold and 2.21-fold observed for the AF11- and AF24-based hexamers, respectively (Figure 5A). However, the neutralization potency of the AF11- and AF24-based hexamers against the B1.351 pseudovirus was substantially enhanced, with improvements of 51-fold and 30-fold, respectively (IC_50_ = 0.8 and 8 pM). Comparable enhancements were observed against the B1.351 virus, with 15-fold and 25-fold increases in potency, respectively (IC_50_ = 2.1 and 6.3 pM). This suggests that the increased neutralization efficiency is likely due to the greater steric hindrance introduced by the Fc-fusion, compared to trimers, which more effectively blocks the interaction with ACE2. These hexamers demonstrated better performance to the monoclonal antibody SA55, which exhibited IC_50_ values of 5.1 pM and 12.2 pM against the B1.351 pseudovirus and virus, respectively. SA55 is a broad-spectrum anti-SARS-CoV-2 antibody that has demonstrated high efficacy against various SARS-CoV-2 variants, including Omicron sublineages [35]. It was identified from a large antibody library derived from SARS convalescents who were vaccinated against SARS-CoV-2. Multimerization can be a way to overcome the decrease of affinity of a monomeric binder induced by mutations associated with viruses variants. AF11- and AF24-based trimers and hexamers were able to neutralize not only B1.351, but also D614G, TY7, BA.1, weakly BA.1 and not EG.5.1.3 and JN1 (Figure 5). It is likely that EG.5.1.3 and JN.1 which are genetically distinct, originating from different lineages, XBB and BA.2, respectively, are both too distant from B1.351 (BA.1 lineage) [39]. According to our models (Figure 6), mutations in the JN1 variant directly impact the epitope recognized by AF11, particularly the K484A substitution. In contrast, substitutions such as K484A, F486P, and Y505H are located at the periphery of the epitope targeted by AF24. These sequence differences, and/or probable structural differences induced, may explain the lack of JN1 neutralization observed for both AF11 and AF24.

Various alternative affinity proteins to conventional IgGs have been developed to neutralize SARS-CoV-2, including VHH fragments [40,41], Affibodies [42], DARPins [43], and αREPs [44]. These efforts resulted in binders with affinities for the RBD ranging from sub-nanomolar to hundreds of nanomolar levels, though their neutralization capabilities were generally limited in their monomeric form. To enhance their neutralization potency, most of these binders - except for Affibodies, which showed IC_50_ values of 1.1 to 2.3 µM - were engineered into dimeric or trimeric formats using peptide linkers, Fc-fusions, or a multimerization domain. These modifications significantly improved their IC_50_ values, achieving as low as 4.0 nM [40], 43 pM [41], 24 pM [43], and 8 nM [44]. Other Affitins were recently engineered into a heterotetramer using a 15-amino-acid linker, achieving an affinity of 90 pM for the RBD and an IC_50_ of 61 pM against the D614G pseudovirus [45]. Each monomeric binding unit of the tetramer exhibited K_D_ values of 380, 31, 15, and 0.8 nM, respectively. Interestingly, our AF11-11-11 Affitin (K_D_ = 22.8 pM, IC_50_ = 41 pM) demonstrates slightly higher potency than this tetramer, despite being a homotrimer composed of monomeric units with a K_D_ of 211 nM for the RBD. This observation suggests that factors beyond the affinity of individual monomers, such as the recognized epitope(s) and the nature or length of the linker, play critical roles in designing highly potent neutralizing multimers. Our pipeline, which incorporates GoldenGate assembly to rapidly generate diverse combinations of monomers, is particularly suited to discovering optimal multimeric constructs. Furthermore, we engineered even more potent hexamers by incorporating Fc-fusions (e.g., (AF11-11-11)_2_-hFc1, K_D_ = 23.4 pM, IC_50_ = 0.8–2.1 pM) ranking them among the most potent neutralizing protein for B.1.351. Indeed, to the best of our knowledge, potent monoclonal antibodies described to neutralize B.1.351 variant have shown IC_50_ from around 1 ng/mL (∼7 pM) [46]. Fc fusions are commonly employed to generate bivalent or bispecific constructs, enhancing therapeutic efficacy by enabling complement activation (CDC) and antibody-dependent cellular cytotoxicity (ADCC). Despite possessing six valencies, potentially with different specificities, the hexameric Affitins fused to Fc maintain a simpler structure and a significantly smaller size (∼102 kDa) compared to mAbs. However, this molecular weight is notably larger than that of monomeric or trimeric Affitins, which may contribute to a prolonged in vivo half-life. Where Fc-mediated functions are not required, future research could explore whether hexameric Affitins assembled solely with peptide linkers or smaller multimerization domains can achieve comparable or even superior efficacy. An Affitin molecule measures approximately 3 x 3 nm, and while multimerization increases its molecular weight, such as in a trimer or in a hexameric Fc, their linear-fusion architecture suggests that these constructs could have a better access to conserved epitopes within deep viral grooves than mAbs which measure approximately 16 x 12 nm.

## 5. Conclusions

Overall, our study highlights the advantages of Affitins making them ideally suited for the development of a streamlined pipeline to produce highly potent neutralizing and multimeric proteins. As a case study, we successfully generated different formats of anti-spike multimers that rank among the most potent neutralizing proteins reported to date. We anticipate that this Affitin-based platform will prove valuable in generating neutralizing proteins against other viruses responsible for infectious diseases.

## Acknowledgements

We thank the P4EU network (Protein Production and Purification Partnership in Europe) and in particular Florian Krammer (Icahn School of Medicine at Mount Sinai, New York, USA) and Sebastiano Pasqualato (European Institute of Oncology, Milan, Italy) for the spike, RBD expression vectors, and Kelvin Lau (Protein production and structure core facility, Lausanne, Switzerland) for ACE2 expression vectors. We thank Hervé Nozach (CEA, France) for the map of the Fc fusion expression vector. We are grateful to Patrick England and Sylviane Hoos (Plate-Forme de Biophysique moléculaire, Institut Pasteur, Paris, France), and to Mike Maillasson (IMPACT Core Facility, BioCore, Nantes Université, France), for the BLI and SPR measurements, respectively. NanoDSF and mass photometry studies were supported by funding from the Biogenouest network, from the “Pays de la Loire” region and from IBiSA. This work was supported by the ANR-20-CO10-0001-01 (COVAFFIT) with financial support for T.M, and by the Région des Pays de la Loire (Trajectoire Nationale). Also funding from Fundação para a Ciência e a Tecnologia (FCT), I.P.; within the scope of the project UIDP/04378/2020 and UIDB/04378/2020 of UCIBIO; the project LA/P/0140/2020 of i4HB; and 2023.10437.CPCA.A2.

## Supplementary data

Supplementary tables S1 to S9, and figures S1 to S8 are available online.

