## Supplemental material for "Optimized pipeline for generating highly potent neutralizing Affitins: application to SARS-CoV-2 spike protein"

**Table S1. Sequences of proteins used in the study.**

| Protein | Sequence | $e_m$ ($M^{-1}cm^{-1}$)* |
|---|---|---|
| RBD (B1.351) | QRVQPTESIVRFPNITNLCPFGEVFNATRFASVYAWNRKRISNCVA<br>DYSVLYNSASFSTFKCYGVSPTKLNDLCFTNVYADSFVIRGDEVR<br>QIAPGQTGNIADYNYKLPDDFTGCVIAWNSNNLDSKVGGNYNYL<br>YRLFRKSNLKPFERDISTEIYQAGSTPCNGVKGFNCYFPLQSYGFQ<br>PTYGVGYQPYRVVVLSFELLHAPATVCGPKKSTNLVKNKCVNFH<br>HHHHHGSGGLNDIFEAQKIEWHE | 40840 |
| spike (B1.351) | QCVNLTTRTQLPPAYTNSFTRGVYYPDKVFRSSVLHSTQDLFLPFF<br>SNVTWFHAIHVSGTNGTKRFDNPVLPFNDGVYFASTEKSNIIRGWI<br>FGTTLDSKTQSLLIVNNATNVVIKVCEFQFCNDPFLGVYYHKNNK<br>SWMESEFRVYSSANNCTFEYVSQPFLMDLEGKQGNFKNLREFVF<br>KNIDGYFKIYSKHTPINLVRDLPQGFSALEPLVDLPIGINITRFQTLL<br>ALHRSYLTPGDSSSGWTAGAAAYYVGYLQPRTFLLKYNENGTIT<br>DAVDCALDPLSETKCTLKSFTVEKGIYQTSNFRVQPTESIVRFPNIT<br>NLCPFGEVFNATRFASVYAWNRKRISNCVADYSVLYNSASFSTFK<br>CYGVSPTKLNDLCFTNVYADSFVIRGDEVRQIAPGQTGNIADYNY<br>KLPDDFTGCVIAWNSNNLDSKVGGNYNYLYRLFRKSNLKPFERDI<br>STEIYQAGSTPCNGVKGFNCYFPLQSYGFQPTYGVGYQPYRVVVL<br>SFELLHAPATVCGPKKSTNLVKNKCVNFNFNGLTGTGVLTESNK<br>KFLPFQQFGRDIADTTDAVRDPQTLEILDITPCSFGGVSVITPGTNT<br>SNQVAVLYQDVNCTEVPVAIHADQLTPTWRVYSTGSNVFQTRAG<br>CLIGAEHVNNSYECDIPIGAGICASYQTQTNSPASVASQSIIAYTMS<br>LGAENSVAYSNNSIAIPTNFTISVTTEILPVSMTKTSVDCTMYICGD<br>STECSNLLLQYGSFCTQLNRALTGIAVEQDKNTQEVFAQVKQIYK<br>TPPIKDFGGFNFSQILPDPSKPSKRSFIEDLLFNKVTLADAGFIKQY<br>GDCLGDIAARDLICAQKFNGLTVLPPLLTDEMIAQYTSALLAGTIT<br>SGWTFGAGAALQIPFAMQMAYRFNGIGVTQNVLYENQKLIANQF<br>NSAIGKIQDSLSSTASALGKLQDVVNQNAQALNTLVKQLSSNFGA<br>ISSVLNDILSRLDPPEAEVQIDRLITGRLQSLQTYVTQQLIRAAEIRA<br>SANLAATKMSECVLGQSKRVDFCGKGYHLMSFPQSAPHGVVFLH<br>VTYVPAQEKNFTTAPAICHDGKAHFPREGVFVSNGTHWFVTQRN<br>FYEPQIITTDNTFVSGNCDVVIGIVNNTVYDPLQPELDSFKEELDK<br>YFKNHTSPDVDLGDISGINASVVNIQKEIDRLNEVAKNLNESLIDL<br>QELGKYEQYIKWPSGRLVPRGSPGSGYIPEAPRDGQAYVRKDGE<br>WVLLSTFLGHHHHHHGSGGLNDIFEAQKIEWHE | 149825 |
| ACE2 | STIEEQAKTFLDKFNHEAEDLFYQSSLASWNYNTNITEENVQNMN<br>NAGDKWSAFLKEQSTLAQMYPLQEIQNLTVKLQLQALQQNGSSV<br>LSEDKSKRLNTILNTMSTIYSTGKVCNPDNPQECLLLEPGLNEIMA<br>NSLDYNERLWAWESWRSEVGKQLRPLYEEYVVLKNEMARANH<br>YEDYGDYWRGDYEVNGVDGYDYSRGQLIEDVEHTFEEIKPLYEH<br>LHAYVRAKLMNAYPSYISPIGCLPAHLLGDMWGRFWTNLYSLTV<br>PFGQKPNIDVTDAMVDQAWDAQRIFKEAEKFFVSVGLPNMTQGF<br>WENSMLTDPGNVQKAVCHPTAWDLGKGDFRILMCTKVTMDDFL<br>TAHHEMGHIQYDMAYAAQPFLLRNGANEGFHEAVGEIMSLSAAT<br>PKHLKSIGLLSPDFQEDNETEINFLLKQALTIVGTLPFTYMLEKWR<br>WMVFKGEIPKDQWMKKWWEMKREIVGVVEPVPHDETYCDPAS<br>LFHVSNDYSFIRYYTRTLYQFQFQEALCQAAKHEGPLHKCDISNS<br>TEAGQKLFNMLRLGKSEPWTLALENVVGAKNMNVRPLLNYFEPL<br>FTWLKDQNKNSFVGWSTDWSPYADGSLEVLFQGPMDEPRGPTIK<br>PCPPCKCPAPNLLGGPSVFIFPPKIKDVLMISLSPIVTCVVVDVSED<br>DPDVQISWFVNNVEVHTAQTQTHREDYNSTLRVVSALPIQHQDW<br>MSGKEFKCKVNNKDLPAPIERTISKPKGSVRAPQVYVLPPPEEEM<br>TKKQVTLTCMVTDFMPEDIYVEWTNNGKTELNYKNTEPVLDSDG<br>SYFMYSKLRVEKKNWVERNSYSCSVVHEGLHNHHTTKSFSRTPG<br>KHHHHHHHHHH | 185025 |

| | | |
|---|---|---|
| AF4 | MRGSHHHHHHGSATKVKFKDYGEEKEVDISKITHVWRVGKMITF<br>GYDDNGKWGTGQVSEKDAPKELLEKLKLN | 13940 |
| AF5 | MRGSHHHHHHGSATKVKFKAPGEEKEVDISKILHVTRVGKMIIFH<br>YDDNGKLGTGGVSEKDAPKELLEKLKLN | 1280 |
| AF7 | MRGSHHHHHHGSATKVKFKVVGEEKEVDISKIYLVRRVGKMILF<br>DYDDNGKYGWGDVSEKDAPKELLEKLKLN | 9530 |
| AF8 | MRGSHHHHHHGSATKVKFKDYGEEKEVDISKILDVNRVGKMIWF<br>FYDDNGKQGVGDVSEKDAPKELLEKLKLN | 8250 |
| AF11 | MRGSHHHHHHGSATKVKFKLYGEEKEVDISKIMDVNRVGKMILF<br>GYDDNGKRGQGWVSEKDAPKELLEKLKLN | 8250 |
| AF15 | MRGSHHHHHHGSATKVKFKYPGEEKEVDISKIVAVLRVGKMIVF<br>DYDDNGKIGGGVVSEKDAPKELLEKLKLN | 2560 |
| AF24 | MRGSHHHHHHGSATKVKFKSDGEEKEVDISKIKHVARGPGASRA<br>LILFHYDDNGKIGTGWVSEKDAPKELLEKLKLN | 6970 |
| AF5-15 | MRGSHHHHHHGSATKVKFKAPGEEKEVDISKILHVTRVGKMIIFH<br>YDDNGKLGTGGVSEKDAPKELLEKLKGPSGQAGAAASESLFVSN<br>HAYGATKVKFKYPGEEKEVDISKIVAVLRVGKMIVFDYDDNGKI<br>GGGVVSEKDAPKELLEKLKLN | 5120 |
| AF5-24-5 | MRGSHHHHHHGSATKVKFKAPGEEKEVDISKILHVTRVGKMIIFH<br>YDDNGKLGTGGVSEKDAPKELLEKLKGPSGQAGAAASESLFVSN<br>HAYGATKVKFKSDGEEKEVDISKIKHVARGPGASRALILFHYDDN<br>GKIGTGWVSEKDAPKELLEKLKGPSGQAGAAASESLFVSNHAYG<br>ATKVKFKAPGEEKEVDISKILHVTRVGKMIIFHYDDNGKLGTGGV<br>SEKDAPKELLEKLKLN | 12090 |
| AF24-5-24 | MRGSHHHHHHGSATKVKFKSDGEEKEVDISKIKHVARGPGASRA<br>LILFHYDDNGKIGTGWVSEKDAPKELLEKLKGPSGQAGAAASESL<br>FVSNHAYGATKVKFKAPGEEKEVDISKILHVTRVGKMIIFHYDDN<br>GKLGTGGVSEKDAPKELLEKLKGPSGQAGAAASESLFVSNHAYG<br>ATKVKFKSDGEEKEVDISKIKHVARGPGASRALILFHYDDNGKIG<br>TGWVSEKDAPKELLEKLKLN | 17780 |
| AF11-11-11 | MRGSHHHHHHGSATKVKFKLYGEEKEVDISKIMDVNRVGKMILF<br>GYDDNGKRGQGWVSEKDAPKELLEKLKGPSGQAGAAASESLFV<br>SNHAYGATKVKFKLYGEEKEVDISKIMDVNRVGKMILFGYDDNG<br>KRGQGWVSEKDAPKELLEKLKGPSGQAGAAASESLFVSNHAYG<br>ATKVKFKLYGEEKEVDISKIMDVNRVGKMILFGYDDNGKRGQG<br>WVSEKDAPKELLEKLKLN | 27310 |
| AF24-24-24 | MRGSHHHHHHGSATKVKFKSDGEEKEVDISKIKHVARGPGASRA<br>LILFHYDDNGKIGTGWVSEKDAPKELLEKLKGPSGQAGAAASESL<br>FVSNHAYGATKVKFKSDGEEKEVDISKIKHVARGPGASRALILFH<br>YDDNGKIGTGWVSEKDAPKELLEKLKGPSGQAGAAASESLFVSN<br>HAYGATKVKFKSDGEEKEVDISKIKHVARGPGASRALILFHYDDN<br>GKIGTGWVSEKDAPKELLEKLKLN | 23470 |
| $(AF11_3)_2$-hFc1 | GSATKVKFKLYGEEKEVDISKIMDVNRVGKMILFGYDDNGKRGQ<br>GWVSEKDAPKELLEKLKGPSGQAGAAASESLFVSNHAYGATKVK<br>FKLYGEEKEVDISKIMDVNRVGKMILFGYDDNGKRGQGWVSEK<br>DAPKELLEKLKGPSGQAGAAASESLFVSNHAYGATKVKFKLYGE<br>EKEVDISKIMDVNRVGKMILFGYDDNGKRGQGWVSEKDAPKELL<br>EKLKLGSSSSDKTHTCPPCPAPELLGGPSVFLFPPKPKDTLMISRTP<br>EVTCVVVDVSHEDPEVKFNWYVDGVEVHNAKTKPREEQYNSTY<br>RVVSVLTVLHQDWLNGKEYKCKVSNKALPAPIEKTISKAKGQPR<br>EPQVYTLPPSRDELTKNQVSLTCLVKGFYPSDIAVEWESNGQPEN<br>NYKTTPPVLDSDGSFFLYSKLTVDKSRWQQGNVFSCSVMHEALH<br>NHYTQKSLSLSPGKGSATKVKFKLYGEEKEVDISKIMDVNRVGK<br>MILFGYDDNGKRGQGWVSEKDAPKELLEKLKGPSGQAGAAASES<br>LFVSNHAYGATKVKFKLYGEEKEVDISKIMDVNRVGKMILFGYD<br>DNGKRGQGWVSEKDAPKELLEKLKGPSGQAGAAASESLFVSNH<br>AYGATKVKFKLYGEEKEVDISKIMDVNRVGKMILFGYDDNGKRG<br>QGWVSEKDAPKELLEKLKLGSSSSDKTHTCPPCPAPELLGGPSVF<br>LFPPKPKDTLMISRTPEVTCVVVDVSHEDPEVKFNWYVDGVEVH<br>NAKTKPREEQYNSTYRVVSVLTVLHQDWLNGKEYKCKVSNKAL | 124620 |

| | | |
|---|---|---|
| | PAPIEKTISKAKGQPREPQVYTLPPSRDELTKNQVSLTCLVKGFYP<br>SDIAVEWESNGQPENNYKTTPPVLDSDGSFFLYSKLTVDKSRWQQ<br>GNVFSCSVMHEALHNHYTQKSLSLSPGK | |
| (AF24$_3$)$_2$-hFc1 | GSATKVKFKSDGEEKEVDISKIKHVARGPGASRALILFHYDDNGK<br>IGTGWVSEKDAPKELLEKLKGPSGQAGAAASESLFVSNHAYGAT<br>KVKFKSDGEEKEVDISKIKHVARGPGASRALILFHYDDNGKIGTG<br>WVSEKDAPKELLEKLKGPSGQAGAAASESLFVSNHAYGATKVKF<br>KSDGEEKEVDISKIKHVARGPGASRALILFHYDDNGKIGTGWVSE<br>KDAPKELLEKLKLGSSSSDKTHTCPPCPAPELLGGPSVFLFPPKPK<br>DTLMISRTPEVTCVVVDVSHEDPEVKFNWYVDGVEVHNAKTKPR<br>EEQYNSTYRVVSVLTVLHQDWLNGKEYKCKVSNKALPAPIEKTIS<br>KAKGQPREPQVYTLPPSRDELTKNQVSLTCLVKGFYPSDIAVEWE<br>SNGQPENNYKTTPPVLDSDGSFFLYSKLTVDKSRWQQGNVFSCSV<br>MHEALHNHYTQKSLSLSPGKGSATKVKFKSDGEEKEVDISKIKHV<br>ARGPGASRALILFHYDDNGKIGTGWVSEKDAPKELLEKLKGPSGQ<br>AGAAASESLFVSNHAYGATKVKFKSDGEEKEVDISKIKHVARGPG<br>ASRALILFHYDDNGKIGTGWVSEKDAPKELLEKLKGPSGQAGAA<br>ASESLFVSNHAYGATKVKFKSDGEEKEVDISKIKHVARGPGASRA<br>LILFHYDDNGKIGTGWVSEKDAPKELLEKLKLGSSSSDKTHTCPP<br>CPAPELLGGPSVFLFPPKPKDTLMISRTPEVTCVVVDVSHEDPEVK<br>FNWYVDGVEVHNAKTKPREEQYNSTYRVVSVLTVLHQDWLNG<br>KEYKCKVSNKALPAPIEKTISKAKGQPREPQVYTLPPSRDELTKNQ<br>VSLTCLVKGFYPSDIAVEWESNGQPENNYKTTPPVLDSDGSFFLY<br>SKLTVDKSRWQQGNVFSCSVMHEALHNHYTQKSLSLSPGK | 116940 |

* Values were obtained using the VectorNTI Advance software (ThermoFisher) and from Expasy website

**Table S2. Primer sequences.**

| | |
|---|---|
| **A. Primers used for silencing BsaI in pFP1001 vector:** | |
| GG-BsaI-del-F | 5'-CTGCAATGATACCGCGAGAACCACGCTCAC-3' |
| GG-BsaI-del-R | 5'-GTGAGCGTGGTTCTCGCGGTATCATTGCAG-3' |
| **B. Primers used for amplification and linearization of pFP1001 vector for GoldenGate assemblies:** | |
| GG-1001-2-F | 5'-GGCTACGGTCTCACTGAAGCTTAATTAATGACTGAGCTTGGAC-3' |
| GG-1001-3-F | 5'-GGCTACGGTCTCTTGAAGCTTAATTAATGACTGAGCTTGGAC-3' |
| GG-1001-R | 5’-GGCTACGGTCTCGATCCGTGATGGTGATGGTG-3’ |
| **C. Primers used for the generation of dimeric and trimeric Affitins:** | |
| GG-AFN.1-F | 5'-GGCTACGGTCTCCGGATCCGCAACAAAAGTAAAGTTC-3' |
| GG-AFN.1-R | 5'-GGCTACGGTCTCCCTTAAGTTTTTCCAGCAGTTCTTTCGG-3' |
| GG-AFN.2-F | 5'-GGCTACGGTCTCCCTACGGTGCAACAAAAGTAAAGTTCAAG-3' |
| GG-AFN.2-R | 5'-GGCTACGGTCTCTTCAGTTTTTCCAGCAGTTCTTTC-3' |
| GG-AFN.3-F | 5'-GGCTACGGTCTCTGCCTATGGTGCAACAAAAGTAAAGTTCAAG-3' |
| GG-AFN.3-R | 5'-GGCTACGGTCTCCTTCAATTTTTCCAGCAGTTCTTTC-3' |
| GG-HMA.1-F | 5'-GGCTACGGTCTCTTAAGGGTCCGAGCGGCCAGGC-3' |
| GG-HMA.1-R | 5’-GGCTACGGTCTCCGTAGGCATGGTTGCTCACAAAC-3’ |
| GG-HMA.2-F | 5’-GGCTACGGTCTCACTGAAGGGTCCGAGCGGCCAGGC-3’ |
| GG-HMA.2-R | 5’-GGCTACGGTCTCTAGGCATGGTTGCTCACAAACAGG-3’ |

**Table S3. Conditions for the ribosome display selection against Spike/RBD.**

| Round | Target presentation (nM when in solution) | Number of washings | Total duration of washings (min) | Number of PCR cycles for RT-PCR |
|---|---|---|---|---|
| 1 | Spike, adsorbed | 6 | 1 | 35 |
| 2 | Spike, adsorbed | 6 | 18 | 25 |
| 3 | RBD-biot, 5 nM | 10 | 23 | 25 |

**Table S4. Binding kinetic parameters for monomers from kinetics assays performed by BLI**

| Affitin | $K_D$ (M) | $k_a$ (1/Ms) | $k_{dis}$ (1/s) | Full $R^2$ |
|---|---|---|---|---|
| **AF4** | $1.56 \times 10^{-8}$ | $4.64 \times 10^{5}$ | $7.25 \times 10^{-3}$ | 0.9541 |
| **AF5** | $3.73 \times 10^{-7}$ | $3.60 \times 10^{5}$ | $1.34 \times 10^{-1}$ | 0.9291 |
| **AF7** | $1.46 \times 10^{-7}$ | $3.53 \times 10^{5}$ | $5.16 \times 10^{-2}$ | 0.9652 |
| **AF8** | $1.12 \times 10^{-7}$ | $5.03 \times 10^{5}$ | $5.64 \times 10^{-2}$ | 0.9447 |
| **AF11** | $2.11 \times 10^{-7}$ | $3.49 \times 10^{5}$ | $7.35 \times 10^{-2}$ | 0.9099 |
| **AF15** | $2.33 \times 10^{-7}$ | $1.78 \times 10^{5}$ | $4.16 \times 10^{-2}$ | 0.9508 |
| **AF24** | $6.03 \times 10^{-7}$ | $1.99 \times 10^{5}$ | $1.20 \times 10^{-1}$ | 0.9486 |

**Table S5. Thermal stability values assessed by NanoDSF for monomeric Affitins and IgG Cv2.1169.**

| Sample ID | Tm1 | | Tm2 | | IP Agg | |
|---|---|---|---|---|---|---|
| | Value | S.D. | Value | S.D. | Value | S.D. |
| **AF4** | ND | NA | ND | NA | ND | NA |
| **AF5** | ND | NA | ND | NA | ND | NA |
| **AF7** | 64.52 | 0.07 | ND | NA | ND | NA |
| **AF8** | 64.66 | 0.98 | ND | NA | ND | NA |
| **AF11** | ND | NA | ND | NA | ND | NA |
| **AF15** | ND | NA | ND | NA | ND | NA |
| **AF24** | 84.68 | 0.46 | ND | NA | ND | NA |
| **IgG Cv2.1169** | 70.23 | 0.02 | 83.03 | 0.13 | 85,22 | 0,30 |

Tm1 and Tm2: thermal transitions midpoints at which 50% of the protein or a domain is denatured. IP Agg: temperature at which 50 % of the protein is aggregated. All proteins were tested at 113 µg/mL. ND = not detected. NA = not applicable.

**Table S6. Binding kinetic parameters for multimers from kinetics assays performed by BLI**

| Affitin | $K_D$ (M) | $k_a$ (1/Ms) | $k_{dis}$ (1/s) | Full $R^2$ |
|---|---|---|---|---|
| **AF 5-15** | $9.52 \times 10^{-9}$ | $1.66 \times 10^{5}$ | 1.58 x 10-3 | 0.9955 |
| **AF 5-24-5** | $4.53 \times 10^{-11}$ | $1.74 \times 10^{6}$ | $7.86 \times 10^{-5}$ | 0.9735 |
| **AF 11-11-11** | $2.28 \times 10^{-11}$ | $1.67 \times 10^{6}$ | $3.79 \times 10^{-5}$ | 0.9807 |
| **AF 24-5-24** | $3.48 \times 10^{-11}$ | $1.34 \times 10^{6}$ | $4.65 \times 10^{-5}$ | 0.983 |
| **AF 24-24-24** | $3.03 \times 10^{-11}$ | $1.45 \times 10^{6}$ | $4.40 \times 10^{-5}$ | 0.9821 |

**Table S7. Thermal stability values of Affitin trimers and hexamers assessed at 100 µg/mL by NanoDSF following freeze/thaw cycles.**

| Sample | Cycle | Tm1 | | Tm2 | | IP Agg | |
|---|---|---|---|---|---|---|---|
| | | Value | S.D. | Value | S.D. | Value | S.D. |
| AF11-11-11 | initial | 75.3 | 1.09 | ND | NA | ND | NA |
| | 1st cycle | 74.57 | 1.41 | ND | NA | ND | NA |
| | 2nd cycle | 73.55 | 0.76 | ND | NA | ND | NA |
| | 3rd cycle | 74.76 | 1.48 | ND | NA | ND | NA |
| | 4th cycle | 73.97 | 0.13 | ND | NA | ND | NA |
| AF24-24-24 | initial | 84.01 | 3.29 | ND | NA | ND | NA |
| | 1st cycle | 85.02 | 1.57 | ND | NA | ND | NA |
| | 2nd cycle | 86.6 | 1.25 | ND | NA | ND | NA |
| | 3rd cycle | 85.85 | 2.04 | ND | NA | ND | NA |
| | 4th cycle | 85.15 | 1.87 | ND | NA | ND | NA |
| $(AF11\text{-}11\text{-}11)_2$-hFc1 | initial | 68,78 | 0,01 | 81,67 | 0,00 | ND | NA |
| | 1st cycle | 68,93 | 0,08 | 81,82 | 0,09 | ND | NA |
| | 2nd cycle | 68,75 | 0,03 | 81,76 | 0,27 | ND | NA |
| | 3rd cycle | 68,98 | 0,08 | 81,65 | 0,04 | ND | NA |
| | 4th cycle | 68,86 | 0,02 | 81,77 | 0,25 | ND | NA |
| $(AF24\text{-}24\text{-}24)_2$-hFc1 | initial | 68,74 | 0,08 | 82,51 | 0,30 | ND | NA |
| | 1st cycle | 68,72 | 0,11 | 82,45 | 0,29 | ND | NA |
| | 2nd cycle | 68,66 | 0,01 | 82,10 | 0,20 | ND | NA |
| | 3rd cycle | 68,64 | 0,02 | 82,06 | 0,11 | ND | NA |
| | 4th cycle | 68,59 | 0,01 | 82,07 | 0,13 | ND | NA |

Tm1 and Tm2: thermal transitions midpoints at which 50% of the protein or a domain is denatured. IP Agg: temperature at which 50 % of the protein is aggregated. All proteins were tested at 113 µg/mL. ND = not detected. NA = not applicable.

**Table S8. Binding kinetic parameters for hFc1 fusions from kinetics assays performed by SPR**

| Affitin | $K_D$ (M) | $k_a1$ (1/Ms) | $K_d1$ (1/s) | $k_a2$ (1/Ms) | $k_d2$ (1/s) | Rmax (RU) | $Chi^2$ ($RU^2$) |
|---|---|---|---|---|---|---|---|
| 11X6-hFc1 | $2.34 \times 10^{-11}$ | $5.19 \times 10^{7}$ | 0.02777 | 0.002184 | $9.97 \times 10^{-5}$ | 538 | 69 |
| 24X6-hFc1 | $1.37 \times 10^{-11}$ | $1.98 \times 10^{7}$ | 0.005711 | 0.00231 | $1.15 \times 10^{-4}$ | 533 | 62 |

**Table S9. Protein contacts and interactions between SARS-CoV 2 spike beta and Affitin 11-11-11 and 24-24-24**

This table is provided as a separate file.

Protein contacts were calculated in Moe with a distance threshold of 4.5Å, and minimum energies for H-Bond, H-Pi and Ionic Bonds of -0.5 kcal/mol. Table abbreviations – Type: D, distance, H, Hydrogen bond, A, Arene, and I, Ionic; Energy : interaction energy in kcal/mol; Distance: distance in Å between the centroids of the interacting atoms ; BB : Denotes whether any of the interacting atoms in the entry are backbone (b) or not (-). The first character represents spike, and the second Affitin.

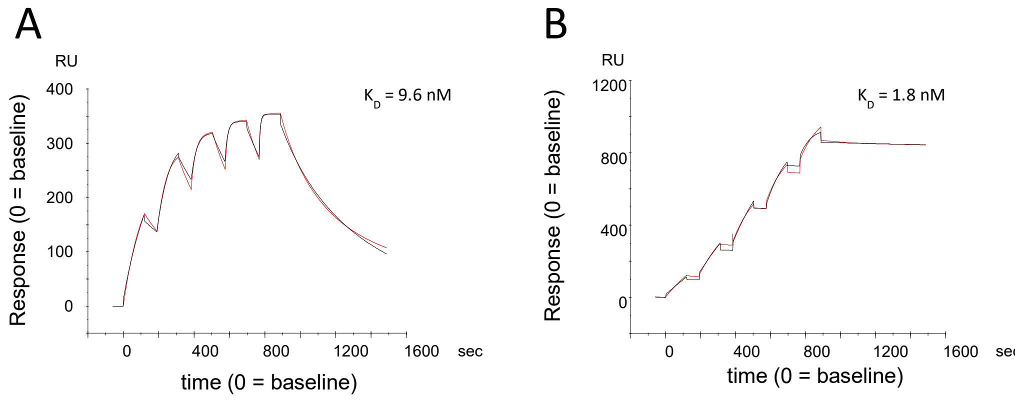


**Figure S1.** Study by SPR of the binding kinetics of RBD (**A**) and spike (**B**) at various concentrations to immobilized ACE-2 on a CM5 chip.

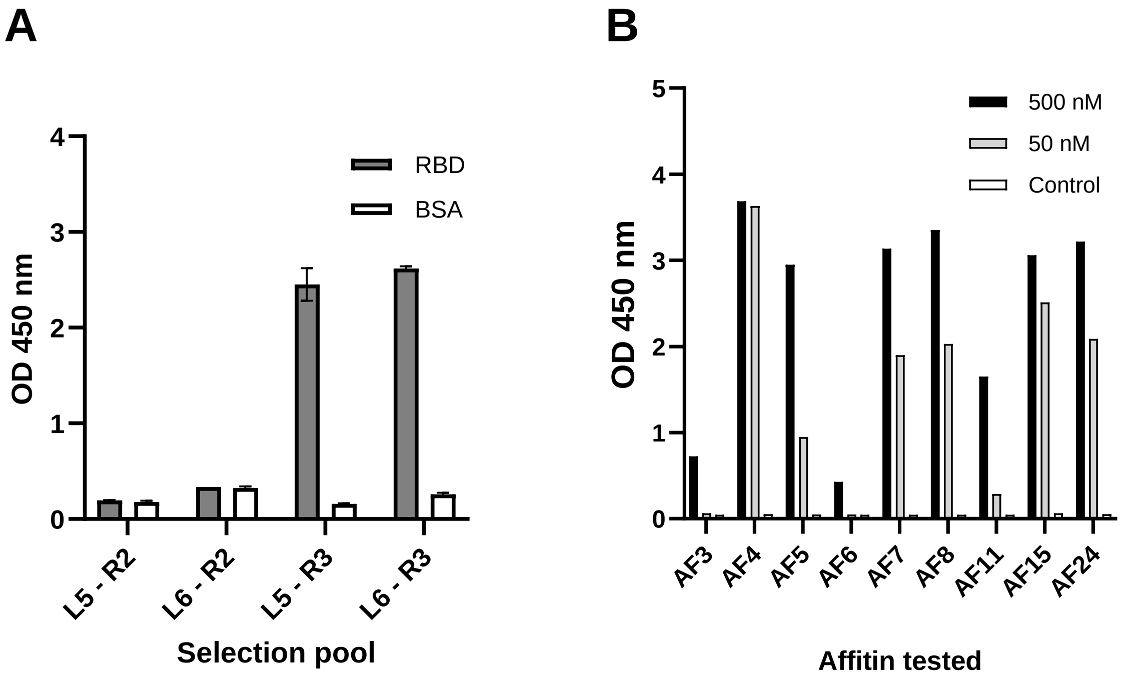


**Figure S2.** Analysis of Affitin selections. (**A**) The output pools of polyclonal Affitins after 2 and 3 rounds of selection (noted R2 and R3) were analyzed via ELISA. In vitro-translated selection pools from libraries L5 and L6 were tested for their binding affinity to RBD immobilized via NeutrAvidin on an ELISA plate. (**B**) Purified monoclonal Affitins were evaluated by ELISA for binding to immobilized RBD at concentrations of 500 nM and 50 nM. For both experiments, controls included wells with only BSA and NeutrAvidin to verify the specificity of the observed binding.

```
L5   ATKVKFKXXGEEKEVDISKIXXVXRVGKMIXFXYDDNGKXGXGXVSEKDAPKELLEKLKLN
AF3  .......AP...........TH.A......I.H......Q.T.A.................
AF5  .......AP...........LH.T......I.H......L.T.G.................
AF6  .......HP...........WN.G......L.E......V.L.G.................
AF15 .......YP...........VA.L......V.D......I.G.V.................
AF7  .......VV...........YL.R......L.D......Y.W.D.................
AF4  .......DY...........TH.W......T.G......W.T.Q.................
AF8  .......DY...........LD.N......W.F......Q.V.D.................
AF11 .......LY...........MD.N......L.G......R.Q.W.................

L6   ATKVKFKXXG EEKEVDISKI KXVXRXXXXX XXXIXFXYDD NGKXGXGXVS EKDAPKELLE KLKLN
AF24 .......SD. .......... .H.A.GPGAS RAL.L.H... ...I.T.W.. .......... .....
```

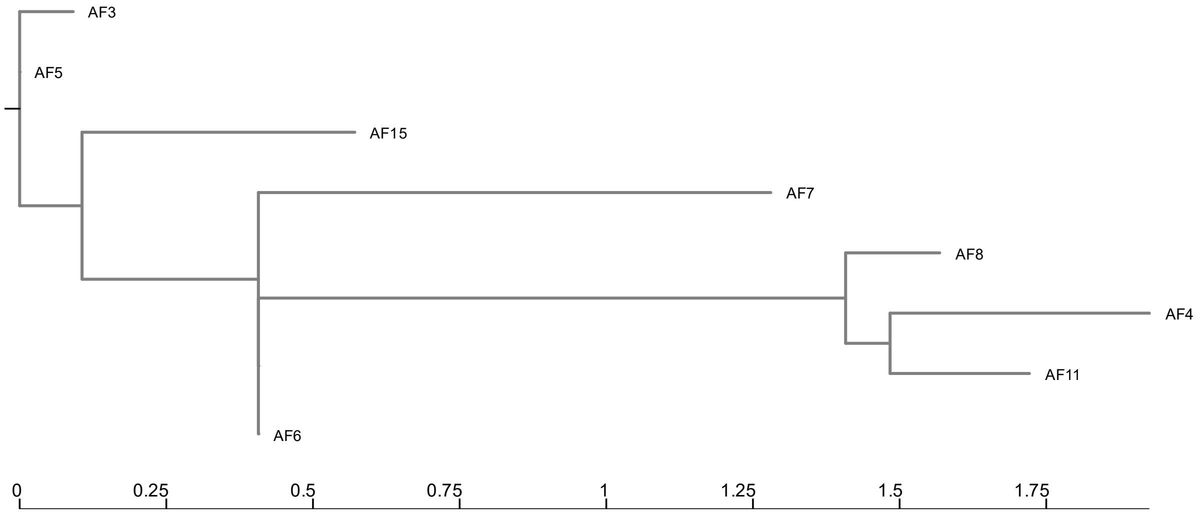


**Figure S3.** Sequences analysis for the anti-RBD Affitins. (**A**) Sequence alignments of designed libraries L5 and L6 and their corresponding isolated Affitins. Randomized residues of designed libraries are represented with a '**X**' and the common residues are indicated by dots. (**B**) A phylogenic tree was created for Affitins from library L5 using ngphylogeny.fr web service in "One click" mode [1]. AF24 was not included in this analysis as it originates from the library L6 and have consequently a quite different sequence than Affitins from the library L5.

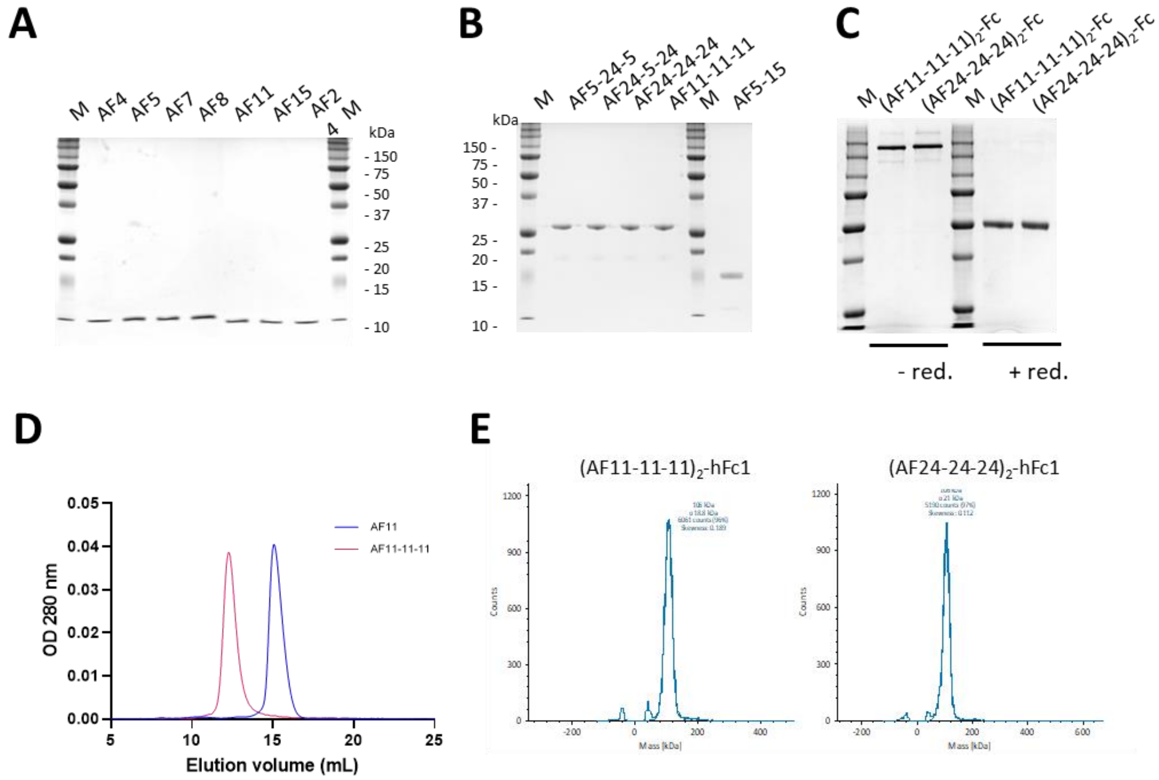


**Figure S4.** Characterizations of anti-RBD Affitins. (**A**) SDS-PAGE analysis of monomers on a 15% gel, (**B**) of the dimeric and trimeric Affitins on a 15% gel; (**C**) of Affitin-Fc fusions on a 12% gel under reducing and non-reducing conditions. (**D**) Typical size-exclusion chromatograms obtained for monomeric and trimeric Affitins eluted from a Superdex 75 column. (**E**) Mass photometry of Affitin-Fc fusions. Gaussians were fitted to the data to estimate the mass and the amount of each specie in the sample.

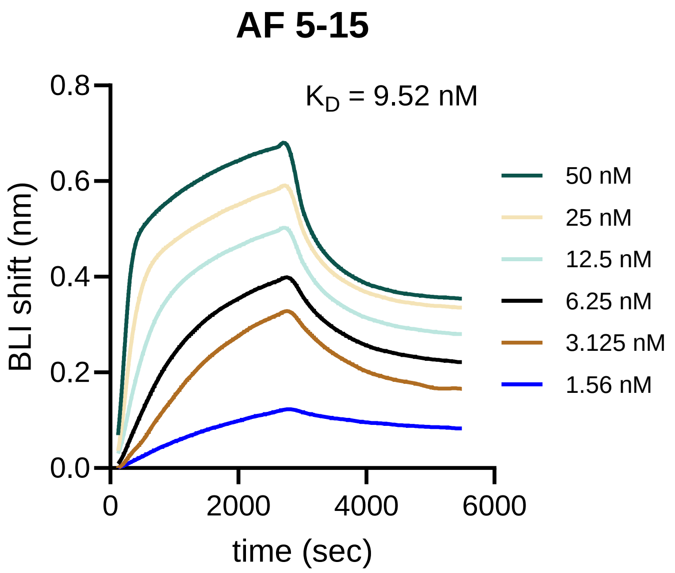


**Figure S5.** Study by BLI of the binding kinetics of the dimer AF 5-15 at various concentrations to RBD immobilized on streptavidin-sensor.

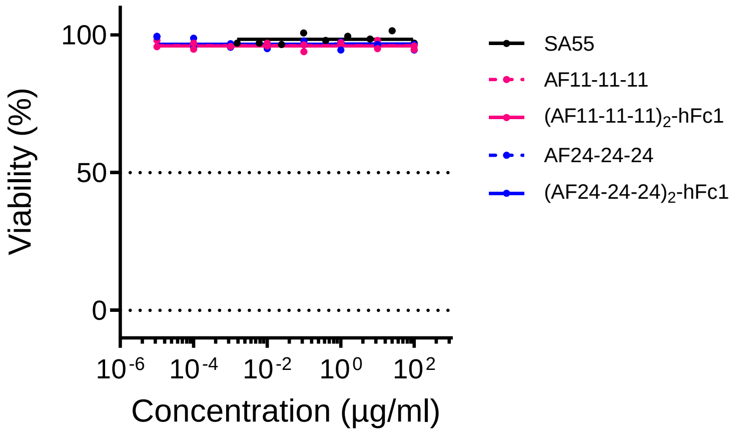


**Figure S6.** Effect of anti-SARS-Cov-2 proteins on the viability of cells used for neutralization tests.

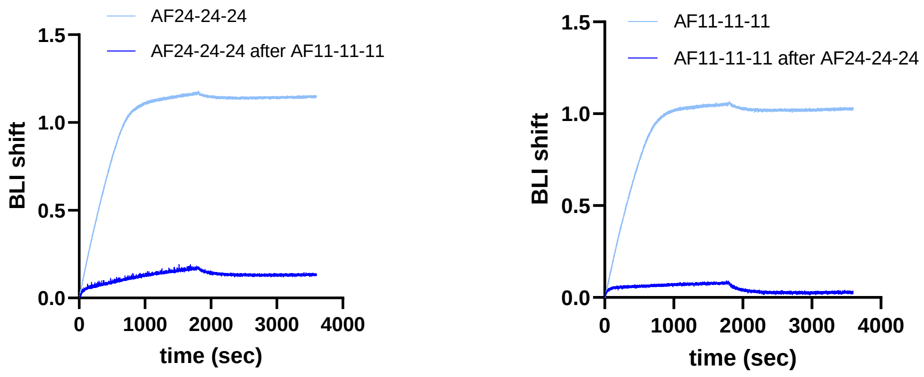


**Figure S7.** Competitive binding assays of trimeric Affitins to immobilized RBD using a streptavidin-coated sensor. (**A**) BLI was used to assess the binding of trimer AF24-24-24, tested alone or following the prior injection of AF11-11-11. (**B**) Similarly, BLI was used to evaluate the binding of trimer AF11-11-11, tested alone or after the prior injection of AF24-24-24.

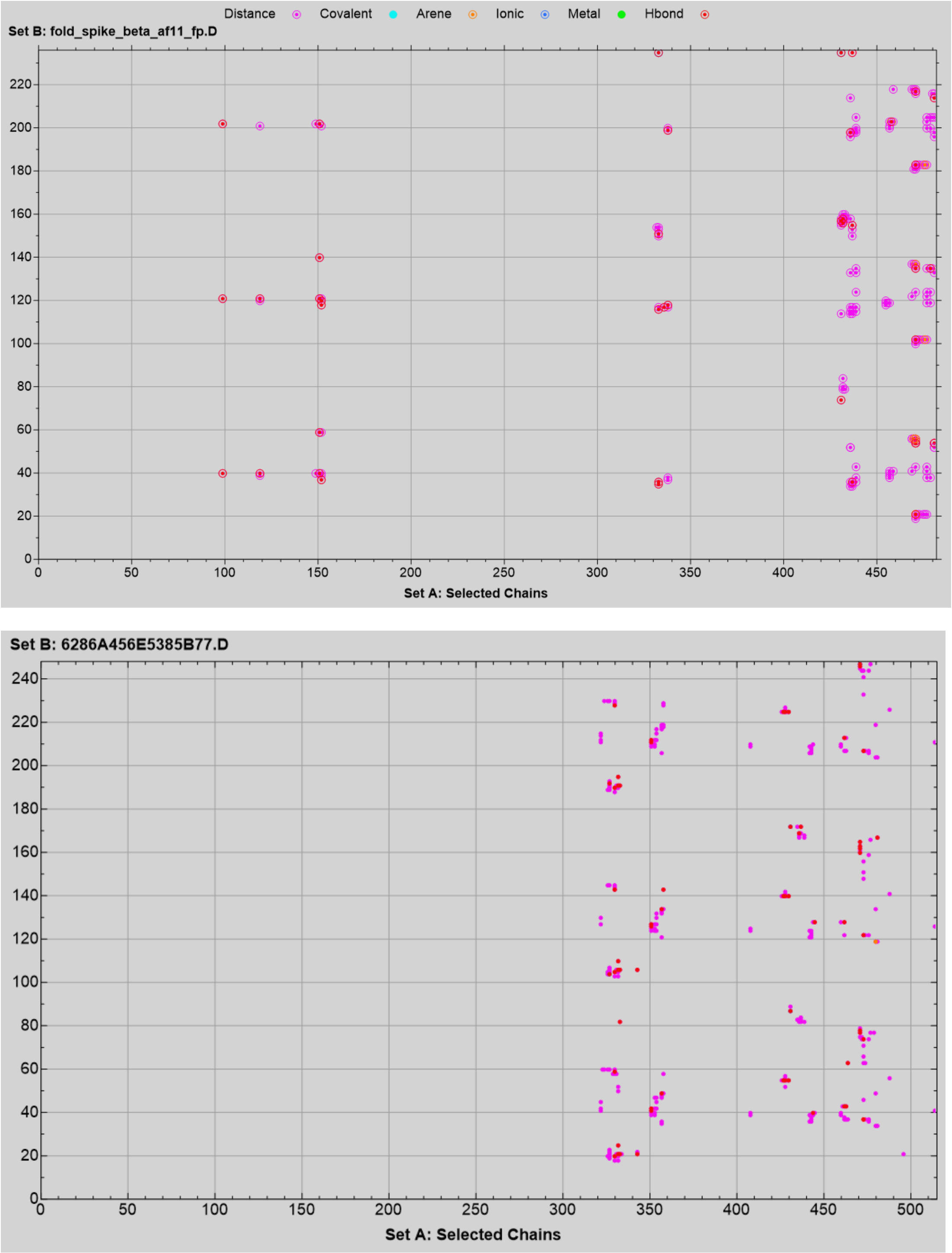


**Figure S8.** Protein-protein interaction contacts plots between spike B1.351 and AF11-11-11 (**A**) and AF24-24-24 (**B**). All contacts have a distance threshold of 4.5Å with hydrogen, aryl-π and ionic bonds with a minimum energy of -0.5 (Table S9).
